# Target-Specificity and Repeatability in Neuro-Cardiac-Guided TMS for Heart-Brain Coupling

**DOI:** 10.1101/2025.02.19.638988

**Authors:** Zi-Jian Feng, Sandra Martin, Ole Numssen, Konstantin Weise, Ying Jing, Gerasimos Gerardos, Carla Martin, Gesa Hartwigsen, Thomas R. Knösche

**Author notes:** Corresponding author, E-mail address (O. Numssen). These authors contributed equally to this work (Zi-Jian Feng, Sandra Martin). These authors contributed equally to this work (Gesa Hartwigsen, Thomas R. Knösche).

## Abstract

The dorsolateral prefrontal cortex (DLPFC) is a principal target for transcranial magnetic stimulation (TMS) in treating major depressive disorder, with therapeutic effects thought to be mediated by its connectivity with the subgenual anterior cingulate cortex. As both regions are involved in autonomic regulation, short-term heart rate changes following DLPFC stimulation may serve as physiological markers to identify stimulation targets. We employed neuro-cardiac guided TMS in a cohort of healthy participants to examine the effects of stimulation intensity and DLPFC target specificity on heart–brain coupling (HBC). We used generalized additive models to assess nonlinear effects of stimulation intensity and target location on HBC, while accounting for pain ratings and other side effects. Intra-subject repeatability across three sessions was evaluated using intraclass correlation coefficients. We observed a non-linear modulation of HBC depending on stimulation intensity and target location, with greater effects at the F3 lateral and F3 posterior targets compared to sham. By evaluating these effects across sessions within participants, we demonstrate the robustness of our results beyond the influence of pain and other side effects on HBC modulation. Exploratory analyses of the directionality show a consistent decrease in HR only at the F3 lateral target with suprathreshold stimulation. These results demonstrate that HBC is modulated in a target- and intensity-specific manner, with particularly consistent effects at F3 lateral sites within the DLPFC. The findings enhance the understanding of TMS-modulated heart-brain interactions, offering a potential framework for optimizing individualized rTMS treatment protocols for depression.

## Introduction

Repetitive transcranial magnetic stimulation (rTMS) is a non-invasive neuromodulation technique widely used to treat psychiatric disorders, particularly major depressive disorder (MDD) [1]. rTMS can induce long-term-potentiation- or long-term-depression-like plasticity, affecting both physiological functions and cognition [2]. Although direct stimulation reaches only circumscribed, superficial cortical areas [3], effects can propagate through downstream connections, impacting remote structures, including subcortical regions [4, 5].

MDD is a network disorder, including a broad dysregulation of the heart-brain axis [6]. A hallmark of this highly-prevalent disorder is reduced vagal activity, leading to elevated heart rate (HR) [7] and reduced heart rate variability [8]. Within this network, the prefrontal cortex is pivotal in cardiovascular autonomic regulation, as demonstrated by lesion and imaging research [9, 10]. Meta-analyses further confirm its essential role in heart rate variability [11].

The frontal–vagal network theory proposes that the dorsolateral prefrontal cortex (DLPFC), subgenual anterior cingulate cortex (sgACC), and the vagus nerve (VN) are central to MDD pathology and autonomic regulation [12]. Clinically, the DLPFC remains a primary rTMS target given its connectivity with the sgACC [13]. Notably, stronger negative DLPFC-sgACC connectivity associates with better therapeutic outcomes [14].

Neuro-Cardiac-Guided TMS (NCG-TMS) was introduced to probe this frontal-vagal pathway [15] using parasympathetic markers, such as HR changes from DLPFC stimulation [16–18], akin to motor evoked potentials elicited by TMS of the primary motor cortex. These measurable, physiological markers are a promising path to optimize therapeutic rTMS applications by individualization of stimulation parameters. Stimulation of the prefrontal cortex, especially of the DLPFC, can lower HR almost immediately [19], likely due to fast conduction via myelinated vagus nerve fibers [12]. Consequently, heart rate may decelerate during rTMS bursts and recover during inter-train intervals [20], reflecting top-down modulation [21, 22]. Although this phenomenon tends to be most pronounced when stimulation engages a DLPFC node within the heart–brain axis, the robustness of the HR modulation [17, 23–25] and its relationship to clinical outcomes remain uncertain [24, 26]. Building on these principles, the NCG-TMS 2.0 protocol [21] aims to identify optimal stimulation targets without reliance on costly functional MRI, using Heart–Brain Coupling (HBC) power as the primary marker of effective engagement of the DLPFC–sgACC–vagus nerve pathway.

While preliminary evidence suggests that NCG-TMS 2.0 can selectively modulate the heart-brain axis and influence heart rate, two key challenges remain: 1) The method exhibits substantial variance, and sample sizes to date have been relatively small, raising concerns about both inter- and intra-subject reliability. 2) HBC quantifies undirected heart rate changes, making it difficult to distinguish genuine neuro-cardiac effects from confounding influences such as pain or discomfort induced by DLPFC stimulation. Furthermore, the lack of publicly available HBC datasets hampers replication efforts and broader validation of the approach.

This preregistered study (https://osf.io/e3fny) systematically evaluates the potential of NCG-TMS 2.0 for DLPFC target site individualization within a rigorous experimental and statistical design. We tested a large healthy cohort across three sessions, comparing six left DLPFC targets alongside a sham condition. We aimed to 1) systematically test whether NCG-TMS 2.0 can reliably modulate heart rate besides effects from stimulation side effects; 2) evaluate repeatability across sessions and generalizability across participants; and 3) determine the optimal NCG-TMS target by assessing a broad range of DLPFC stimulation sites. Additionally, we openly share the acquired dataset to enhance transparency and support future research on this important topic. The results of this study enhance the understanding of rTMS-modulated heart-brain interactions, advancing target selection and individualized stimulation protocols.

## Materials and Methods

### Participants

We included 19 healthy, right-handed participants (9 females, 18-39 years, mean age 30.3 ± 5.5 years). The sample size was based on an a priori power analysis (see Supplementary Methods). Participants were eligible for TMS and inclusion followed the safety guidelines for TMS studies [27]. The experiment complied with the Declaration of Helsinki and was approved by the local ethics committee. Participants provided written informed consent.

### Study Design

The study design comprised a preparatory (Session 0) and an experimental phase (Sessions 1 to 3), involving single-blinded TMS sessions (Fig. 1a). In Session 0, participants underwent an MRI scan to acquire structural images (see Supplementary Methods for details), used for TMS neuronavigation. Subsequently, individual motor thresholds were determined. During sessions 1 to 3, we applied the 10-Hz dash rTMS protocol from NCG-TMS 2.0 [21] to the DLPFC targets identified in Session 0. Sessions were separated by at least one week (mean distance 10.8 ± 9.5 days). The stimulation protocol consisted of 16 blocks, with 15 active trains per target. Intensity increased in 2% increments of the maximum stimulator output (MSO). Each block included a 10 Hz TMS train lasting 5 seconds, followed by an 11-second inter-trial interval (Fig. 1b). The final intensity was set at 120% of the participant’s resting motor threshold (rMT), while the first active intensity was set at 28% MSO below this final level (Fig. 1b). Prior to the active stimulation, a rest block at 0% MSO was included to establish a baseline.

**Figure 1.**
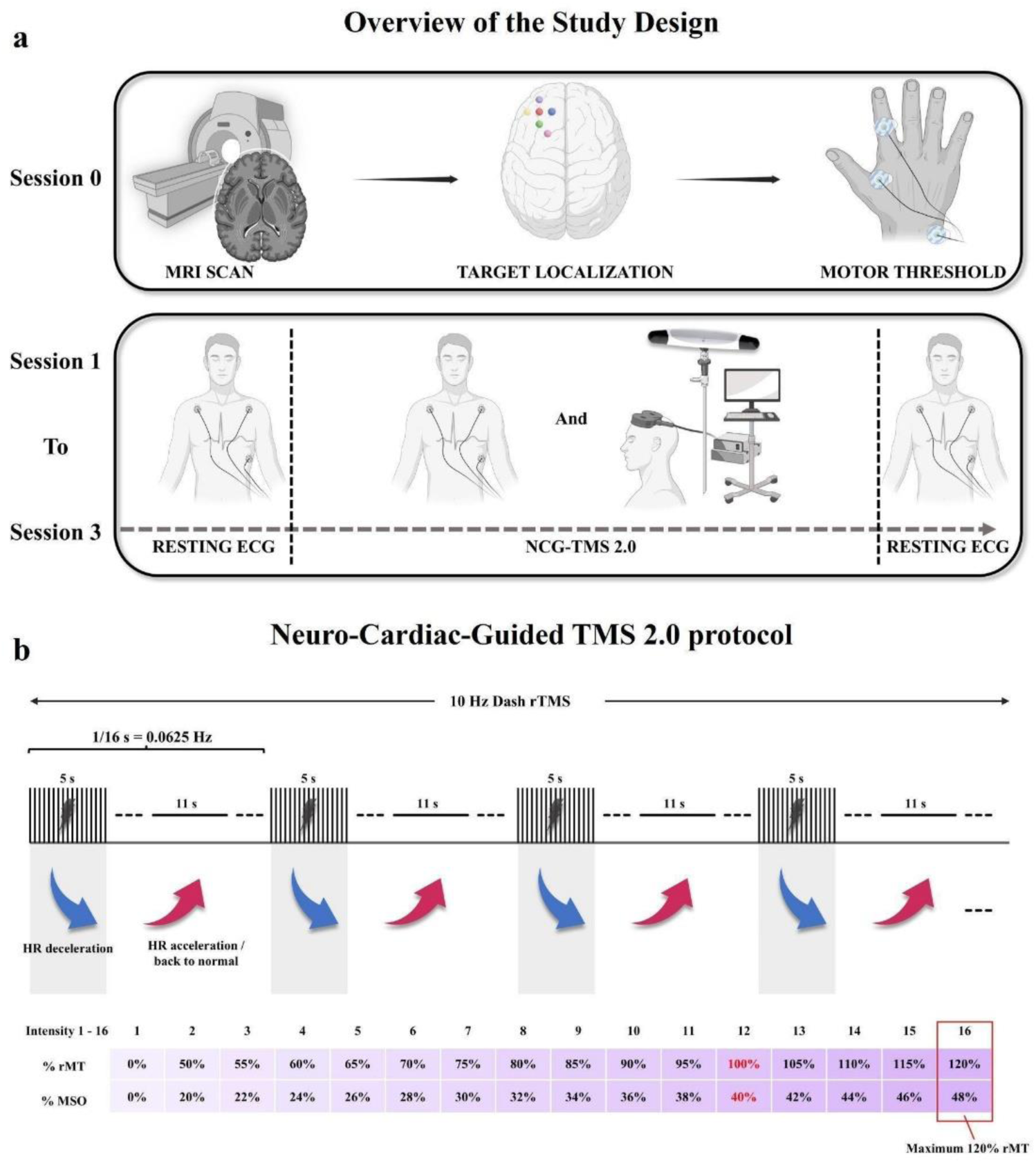
Overview of Study Design and Neuro-Cardiac-Guided TMS 2.0 Protocol. **Panel a** shows the study design. The study consisted of a preparatory (Session 0) and three experimental sessions (Sessions 1–3). During Session 0, anatomical targets within the DLPFC were determined using individual MRI scans, and personalized TMS parameters were established by identifying motor thresholds. Target localization is illustrated by colored circles: F3 (red), F3 lateral (yellow), F3 medial (blue), F3 anterior (purple), F3 posterior (green), and the 5-cm method (pink). Sessions 1 to 3 comprised baseline and post-stimulation resting electrocardiogram (ECG) measurements and the administration of NCG-TMS. **Panel b** shows the specifics of the applied NCG-TMS 2.0 protocol which employs 10 Hz dash rTMS. The protocol involves repeating 16-second cycles, each consisting of a 5-second train of 10 Hz rTMS followed by an 11-second rest interval, yielding an overall entrainment frequency of 0.0625 Hz (1 cycle per 16 seconds). The intensity of stimulation gradually increases through 15 incremental steps (each incremented by 2% of the MSO) and culminating at 120% rMT. An exemplary intensity scheme is illustrated for a participant whose rMT was identified as 40% MSO; thus, the maximum stimulation intensity (120% rMT) corresponds to 48% MSO. Physiological effects on heart rate (HR) during the protocol cycles—characterized by brief deceleration during stimulation followed by acceleration or normalization during rest intervals—are schematically indicated. HR, heart rate; rMT, resting motor threshold; MSO, maximum stimulator output.

Our study incorporated specific adjustments to the original NCG-TMS 2.0 protocol by adding resting electrocardiogram (ECG) periods for baseline correction. Specifically, we introduced eight blocks of rest prior to the initiation of the active stimulation. Additionally, two blocks of post-stimulation resting periods were added to mitigate edge effects when calculating HBC power (Fig. 1a). We stimulated six cortical regions within the left DLPFC to investigate the spatial specificity of NCG-TMS. The precise definitions of these target locations, labeled Beam F3 (short: F3), F3 anterior, F3 posterior, F3 lateral, F3 medial, and 5-cm rule method (short: 5cm), are shown in Figure 1a. Target localization was guided by individual anatomical MRI and the Beam F3 algorithm [21, 28]. Two of the targets (F3 and 5cm) have been investigated previously [21] and four additional targets were defined surrounding the F3 location, positioned 2 cm in the anterior, posterior, lateral, and medial direction. The 5cm target was defined as the stimulation site where the TMS coil was positioned 5 cm anterior to the optimal scalp location for eliciting activation of the FDI muscle [21, 29]. If the 5cm target was within 1-cm of any other target, it was excluded.

For all targets, the coil was oriented 45° relative to the parasagittal plane, with the handle pointing posteriorly. Prior to the main experiment, tolerance was assessed by applying a single 10 Hz TMS train at 120% of the resting motor threshold (rMT) to each target. Discomfort was quantified using the Discrete Visual Pain Rating Scale (DVPRS) [30] with pain scores for each target provided in Figure S1. If a pain score exceeded 6/10 (i.e., above “moderate”), the coil angle was adjusted from 45° to 90° to reduce discomfort (see Supplementary Methods for details). TMS coil positioning was guided by a neuronavigation system to ensure consistent targeting throughout the experiment (see Supplementary Methods for details).

Sham stimulation was performed at one of the five targets, randomly assigned for each participant. Sessions 2 and 3 consisted of repetitions of both active and sham NCG-TMS 2.0 protocols conducted during Session 1. Target order was randomized across participants and kept constant across sessions.

Following each session, a stimulation-related side-effects checklist (Table S1) was administered to systematically assess the severity of 20 symptoms on a 0–4 scale [31]. For each session and target, individual sum scores of side effects and pain ratings from the DVPRS (Fig. S1) were calculated. Table S2 and Figures S2 and S3 summarize the frequency of individual side effects, and the complete measurement procedure is detailed in the Supplementary Methods.

### Data Analysis

#### Individual Heart-Brain Coupling

The ECG preprocessing and HBC analysis procedures were adapted from previously established methods [21]. HBC was quantified by the power at 0.0625 Hz corresponding to one TMS cycle comprising 5 seconds of stimulation and 11 seconds of rest. Details of the HBC computation are provided in the Supplementary Methods. The analysis was applied to all 26 blocks of ECG data. However, only the 16 blocks corresponding to the NCG-TMS 2.0 protocol were retained, while the initial 8 resting blocks and the last 2 resting blocks were excluded to mitigate edge effects. All analyses were conducted using the raw values of HBC power. The high frequency resolution HBC value for each block was then calculated and used for subsequent analyses. Plots for each target across all three sessions for each participant are provided in the supplementary materials (Fig. S4).

### Mixed-Effects Regression Analysis

To analyze the relationship between HBC power of each block and our experimental factors (stimulation intensity and target location) as well as nuisance variables (pain and other side effects), we used a generalized additive mixed-effects model (GAM) since raw data suggested a non-linear relationship between stimulation intensity and HBC (Fig. S5). The GAM was fitted with a Gamma distribution and a log link function to account for the non-normal distribution of HBC power (EQ 1). We also fitted a standard linear regression model to confirm our results without relaxing the linearity assumption. To this end, a generalized linear mixed-effects model (GLMM) was set up similar to the GAM. Further details on the statistical analysis are given in the Supplementary Methods.

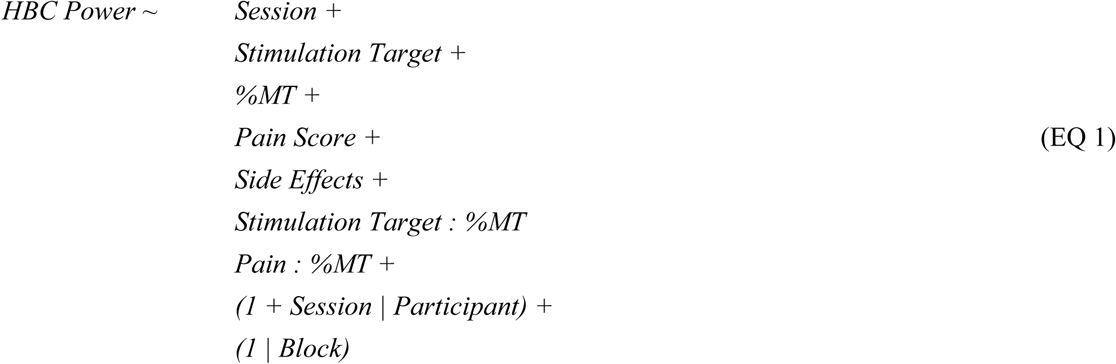

In all regression analyses, we employed simple contrast coding for categorical factors and mean-centering for continuous predictors to ensure interpretability and comparability of the results. Additionally, we applied a false discovery rate (FDR) correction to the p-values obtained from the model summaries to control for multiple comparisons. Analyses were carried out in R version 4.4.0 (R Core Team, 2023) using the packages lme4, mgcv, modelbased, ggeffects, performance, and sjPlot [32–37].

### Repeatability Analysis

To assess the repeatability of block-level measurements across multiple sessions for each target, we calculated the Intraclass Correlation Coefficient (ICC) using a two-way random-effects model. ICCs were computed for each stimulation target to assess the consistency of block-level HBC values across three experimental sessions. Additionally, pairwise ICCs between individual sessions were examined to further explore intra-subject reliability. Details of the ICC computation are provided in the Supplementary Methods.

### Analysis of Heart Rate Modulation Direction

To explore the directionality of stimulation-induced autonomic effects, HR was analyzed within a 2-second window following each TMS train, in line with prior findings on rapid vagal and pupil responses [12, 34]. Pre-train HR was assessed using two reference intervals: 1) a stable baseline (excluding the first two blocks of each session), and 2) the 2-second period immediately prior to stimulation onset (see Supplementary Methods).

## Results

We report data from 19 healthy adults who each completed one preparatory and three experimental sessions. In each session, we applied the NCG-TMS 2.0 protocol [21] to six active targets in the DLPFC and one sham condition, using individual motor thresholds to calibrate stimulation intensity. We refined the NCG-TMS 2.0 protocol by assessing two resting electrocardiogram (ECG) periods (before and after stimulation) for each target to accurately capture stimulation-induced changes to HBC. First, HBC was quantified for each target following the original analysis [21] with an additional baseline correction (see Fig. S4 for individual raw HBC). We then used linear and non-linear mixed-effects regression to analyze the relationship between HBC power of each stimulation block and our experimental factors (stimulation intensity and target location) as well as nuisance variables (pain and other side effects). Moreover, we tested the robustness of our findings by investigating the repeatability of TMS-induced changes in HBC across the three sessions. Finally, we analyzed the direction of HR modulations to pinpoint the underlying modulation mechanisms of the heart-brain axis.

### Target- and Intensity-Specific Modulation of Heart-Brain Coupling

The raw target- and session-wise data suggest a non-linear relationship between stimulation intensity and HBC power (Fig. S5). To account for this non-linearity, we set up a generalized additive mixed-effects model (GAM). We also confirmed our analyses using a standard, generalized linear mixed-effects model (GLMM). We aimed to test 1) whether there is a significant intensity- and target-dependent modulation of HR by TMS, 2) to what extent this effect can be explained by pain or other side effects, and 3) whether the TMS intensity effect on HBC is target-specific. To this end, we tested how well HBC is explained by the experimental variables *stimulation target*, *session*, and *intensity (%MT)*, as well as the covariates *pain* and *side effects*.

#### Effects of Session and Target

The GAM included parametric terms for session number and stimulation targets (6 active targets and 1 sham target). Results revealed no differences in HBC of session 2 and 3 relative to session 1 (β_session 2_ = 1.13, p = 0.810, β_session 3_ = 1.02, p = 0.981). Stimulation targets at F3 anterior, F3 lateral, and F3 posterior were associated with increased HBC relative to sham, while all other targets were linked with reduced HBC compared to sham (Table 1). After FDR correction, this main effect remained significant for the lateral and posterior F3 targets (β_F3lat_ = 1.46, p < 0.001, β_F3pos_ = 1.18, p = 0.033). In sum, there was no effect of session but an effect of target with some stimulation targets showing significantly higher HBC relative to sham.

**Table 1.**
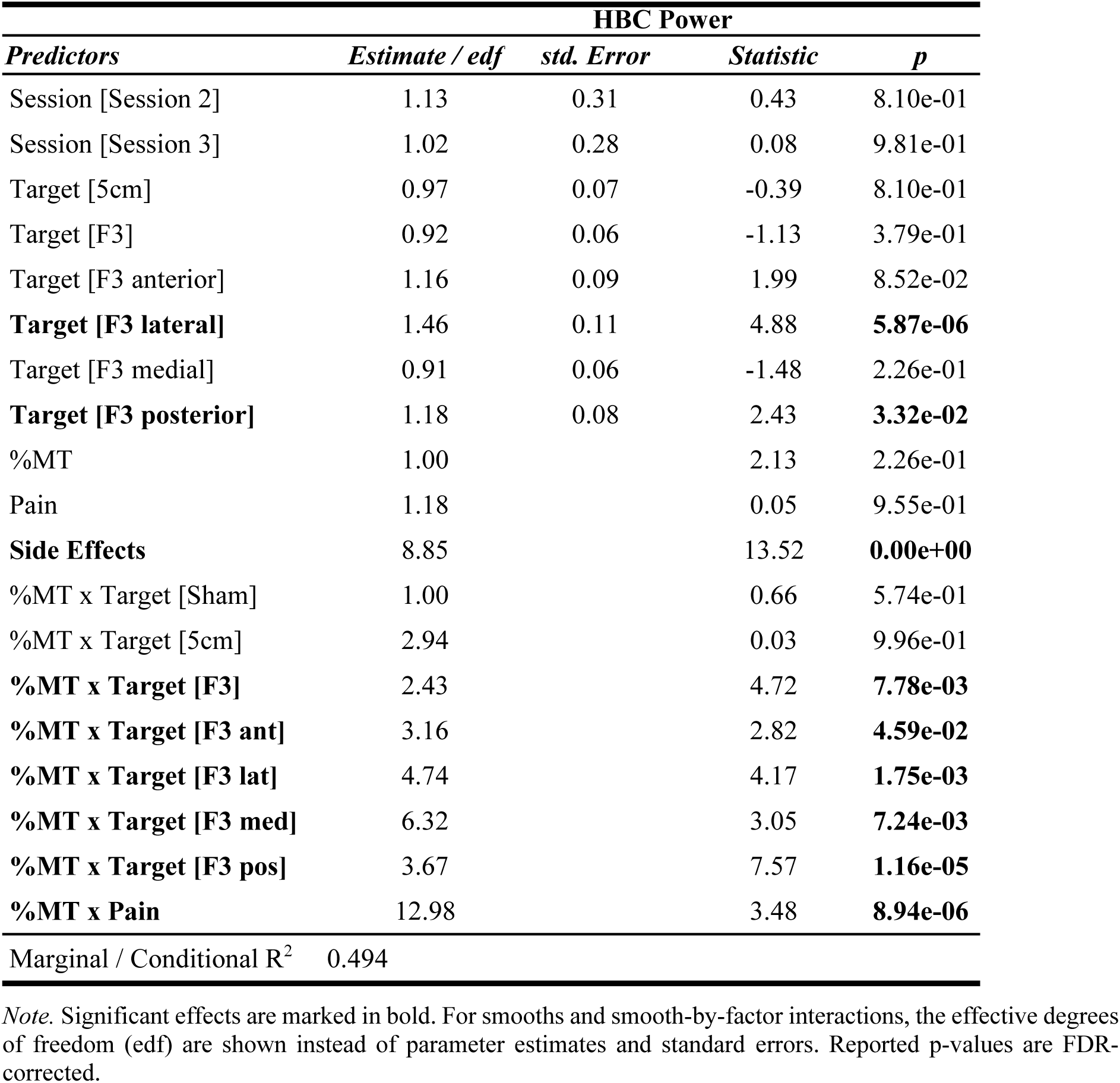
Results from the Generalized Additive Model (GAM).

#### Effects of Stimulation Intensity, Pain, and Side Effects

The GAM further included smoothing splines for %MT, Pain, and a sum score of Side Effects. Results showed a significant effect on HBC only for side effects (F_Side Effects_ = 13.52, p < 0.001, Table 1). Exploring the smoothing curve of Side Effects revealed an inverted U shape, where for low to moderate values (side effects score ≤ 7.33), HBC was positively correlated with the side effects score, and for moderate to high values (side effects score ≥ 7.33), HBC decreased (Fig. S6) with larger scores. In sum, we could not prove main effects for %MT and Pain, but for Side Effects, which showed a non-linear relationship with HBC.

#### Interaction of Stimulation Intensity and Target

Modeling the interaction between %MT and stimulation target showed significant non-linear effects for F3 and surrounding targets (F3_ant_, F3_pos_, F3_med_, F3_lat_; all p < 0.05, FDR-corrected, see Table 1), but not for sham stimulation and the 5cm target (Fig. 2a). To assess the magnitude of target-specific effects, we calculated the difference in HBC power for each target relative to Sham for 60 and 100% MT and averaged across all intensities. Averaging across all stimulation intensities, we found significantly increased HBC for F3 and all surrounding targets (all p < 0.05, FDR-corrected, Fig. 2b). The strongest increase in HBC relative to sham was observed in the posterior, medial, and lateral F3 targets. For example, the posterior F3 target increased HBC by 140 points or 11,145% compared to sham TMS (estimated marginal mean F3_pos_: 138.98, sham: −1.26 points in HBC power). The intensity-dependency of this effect was further confirmed when comparing differences in HBC at lower and higher intensities. While there was no significant modulation of HBC at 60% MT, 100% MT induced increases in HBC in several F3 targets compared to sham. In summary, increasing HBC was modulated by the interaction of %MT and targets, showing that F3 and surrounding targets yield higher HBC in an intensity-dependent manner.

**Figure 2.**
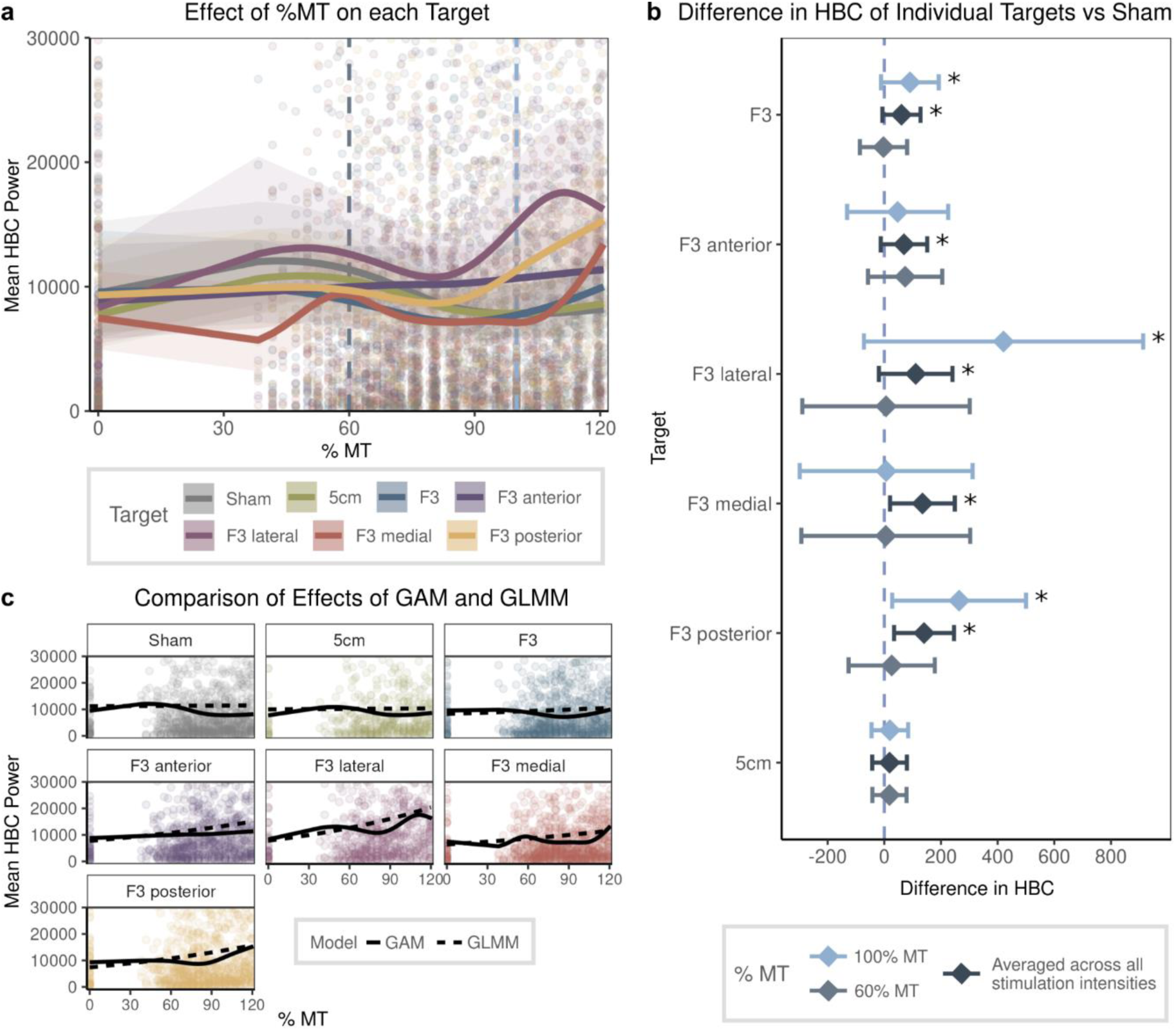
TMS-induced Heart-Brain-Coupling across Different DLPFC Stimulation Sites. **Panel a** displays estimated marginal predictions from a generalized additive model (GAM) for the interaction between stimulation intensity (% motor threshold; % MT) and stimulation target with raw data plotted in the background. Dashed lines show stimulation intensities, for which differences relative to sham were computed (next panel). **Panel b** illustrates the difference in marginal predictions for different values of % MT per target relative to sham. Significant effects for stimulation intensity at 60% MT and averaged across all stimulation intensities are marked with an asterisk. Stimulation over all F3 targets significantly increased HBC power compared to sham. **Panel c** compares marginal predictions for each target derived from the GAM and the generalized linear mixed-effects model (GLMM). Although both models lead to similar conclusions, the GAM outperformed the GLMM in measures of goodness of fit.

#### Interaction of %MT and Pain

Results from the GAM revealed a significant non-linear interaction of %MT and pain (F = 3.48, p < 0.001). To explore this interaction of both continuous predictors, we ran a simple slopes analysis and calculated Johnson-Neyman intervals. Results showed that pain significantly influenced the effect of %MT for low pain values (pain score between 0.67 and 1.33 points; range 0-6) and high pain values (pain scores above 4). For these values, the slope of %MT was positive, thus increasing HBC when pain scores were high (Fig. S7). To further evaluate the contribution of pain to changes in HBC, we set up two additional models: a GAM without the effect of *%MT*, only including *pain* and its interaction with *stimulation target*, and a GAM without *pain scores* but including *%MT* and its interaction with *target*. Comparing AIC and R^2^ values for these models with our final regression model (see EQ 1) showed that the GAM including both terms (*%MT* and *pain*) outperformed the individual models in terms of AIC (see Table S2). In summary, while pain influences HBC, it does not fully account for the %MT dependence. Moreover, %MT seems to be a stronger predictor than pain alone.

#### Comparison of non-linear and linear mixed-effects regression

The GAM demonstrated a superior fit as indicated by a lower Akaike Information Criterion (AIC, AIC_GAM_ = 116,683.0, df_GAM_ = 108.42, AIC_GLMM_ = 117,027.9, df_GLMM_ = 27.0). Results from the GLMM, which corroborated findings from the GAM (Fig. 2c), are presented in Table S3.

#### Interaction of different side effects and stimulation targets

To explore whether certain side effects had a particularly strong influence and if this might be target-specific, we calculated an additional GAM using the three side effects with highest ratings (Muscle contraction, Scalp pain, and Headache, see Table S2) and their interaction with stimulation target as individual predictors instead of the side effect sum score. Results showed interactions across different targets with different side effects (Table S5). Importantly, the interactions of stimulation intensity and targets as described above from our initial GAM remained robust.

For muscle contractions, there was a significant interaction with the targets sham and 5cm, both decreasing HBC, as well as F3 anterior and posterior, both increasing HBC with higher scores. For scalp pain, targets 5cm, F3, F3 anterior, and F3 medial were associated with increasing HBC with higher scores. For the side effect headache, there was a significant interaction with the targets sham and F3 anterior, which were linked to increasing HBC with higher scores, whereas the targets 5cm and F3 medial were linked to decreasing HBC with higher scores. Figure S8 displays the average marginal means per target for each of the three side effects.

### Robustness of Changes in Heart-Brain Coupling across Sessions

We calculated the intraclass correlation coefficient of block-level HBC power to assess the repeatability of block-level measurements across three sessions for each stimulation target (Fig. 3a). Although most targets showed considerable variability (ICC below 0.4), the anterior and lateral F3 targets demonstrated moderate to high repeatability of findings, especially at higher stimulation intensities. To account for a potentially stronger session effect in session 1, we focused on the anterior and lateral F3 targets and explored their pairwise repeatability between sessions. Both targets demonstrated high ICC values between sessions 2 and 3, particularly at higher stimulation intensities, indicating excellent repeatability (Fig. 3b). These results demonstrate the robustness of our findings and further corroborate the findings from our regression analysis, indicating reliable HBC modulation through rTMS over anterior and lateral F3 targets.

**Figure 3.**
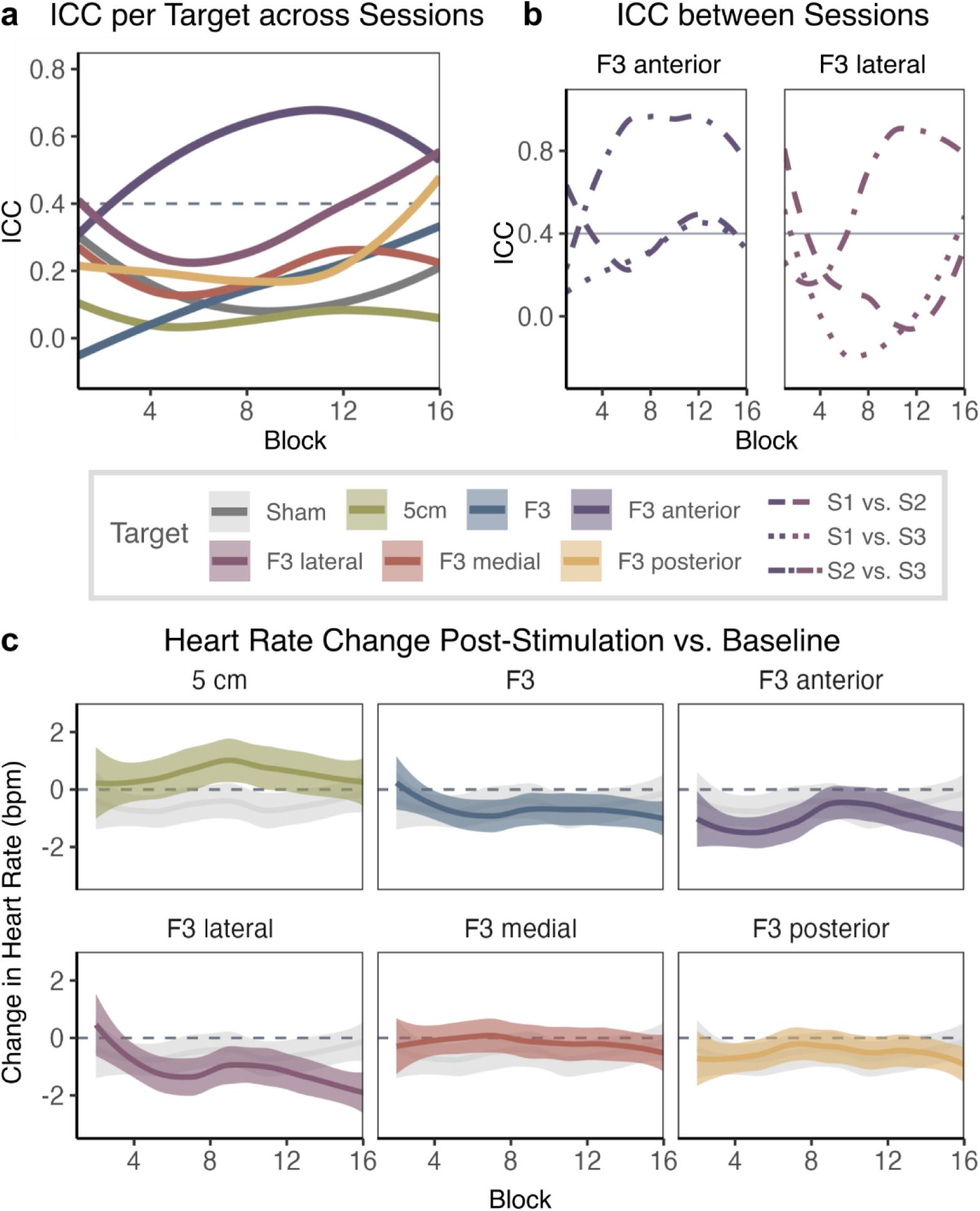
HBC repeatability and Changes in Heart Rate for Block-level Measurements. **Panel a** shows ICC of block-level HBC power (i.e., the spectral power at the 0.0625-Hz entrainment frequency of the NCG-TMS 2.0 protocol) across the three sessions for each stimulation target. An ICC value of 0.4 is considered moderate repeatability (the horizontal dashed line). **Panel b** shows pairwise ICC of block-level HBC power between session pairs (S1–S2, S1–S3, S2–S3) for F3 anterior and F3 lateral which demonstrated smallest variability in total. For both targets, sessions 2 and 3 show highly robust findings. Block numbers on the x-axis correspond to increasing stimulation intensities. **Panel c** shows heart rate changes in beats per minute (bpm) for heart rate recorded post-stimulation versus pre-stimulation baseline for each target. Each target panel shows heart rate change for sham stimulation in grey in the background. Shaded areas indicate 95% confidence intervals.

### TMS-Induced Directionality of Heart Rate Modulation

We used the ECG data before and after rTMS to each target to assess the directionality of TMS-induced changes in HR. Across all targets and sessions, HR fluctuations remained relatively small, with changes typically within ±2 bpm compared to the HR at baseline (Fig. 3c). Notably, high-intensity F3 lateral stimulation was the only condition that consistently and reliably reduced HR with increasing stimulation intensity. Again, this pattern was particularly stable for sessions 2 and 3 (Fig. S9), confirming the reduced variability in stimulation-induced effects in these sessions. Similar directions in HR change were observed in the results for post-train HR changes compared to pre-train HR measurements (Fig. S10–S11).

## Discussion

The NCG-TMS 2.0 protocol was developed to quantify TMS-induced heart rate changes via the DLPFC–sgACC–vagus nerve pathway [21], offering a potentially cost-effective and easy-to-implement approach to individualize MDD treatment with rTMS. Here, we systematically evaluated the protocol’s effectiveness and robustness in enhancing HBC across a set of DLPFC targets, to identify the optimal site to target the heart-brain axis. Our findings show that specific DLPFC locations––notably F3 lateral and posterior––were associated with a systematic significant increase in HBC relative to sham. Across sessions, TMS-induced heart-rate modulation was particularly robust for the lateral F3 target. Importantly, although pain and side effects significantly influenced HBC, they did not fully account for the stimulation-induced changes, indicating a causal effect of rTMS on the heart-brain axis.

Our findings reveal the crucial role of stimulation site and stimulation intensity to reliably modulate HBC. They demonstrate a non-linear and target-specific effect of stimulation intensity on HBC, which partially aligns with previous findings, which observed a (linear) main effect of intensity but lacked a sham condition to demonstrate target specificity [21]. A recent study using sgACC-guided rTMS similarly demonstrated significant HBC modulation in targets around F3, though generalizability was limited by spatial heterogeneity and small sample size [38]. Importantly, we critically put the NCG protocol to the test by further evaluating the directionality of TMS-induced HBC, providing further insight into the underlying neurobiological mechanisms. Due to its nature, the HBC metric does not differentiate between the directionality of the induced heart rate change, i.e., between heart rate increases or decreases. Our results suggest that especially high-intensity F3 lateral stimulation reliably reduces heat rate. A meta-analysis of NCG-TMS research demonstrated that stimulation at frontal regions (F3/F4, FC3/FC4) generally lowers heart rate, whereas motor cortex stimulation (C3/C4) typically elevates heart rate [12]. By contrast, more recent findings suggest that stimulation at C3/C4 can also reduce heart rate, with no significant difference observed compared to frontal stimulation (F3/FC3) when targeting the left hemisphere [16]. This implies that the motor cortex may not be an ideal control target due to confounding heart rate effects. In our study, sham rTMS mitigated these confounds and improved comparability. Notably, the 5cm target did not significantly alter HBC, differing from prior findings of strong individual responses at that location [21, 38].

Stimulation intensity is a critical parameter influencing the effect of TMS [39, 40]. While subthreshold rTMS has been shown to induce structural synaptic plasticity in animal models [41] and modulate cognitive processes in humans [42], its influence on heart rate or other heart-brain axis-related states remains inconclusive [43]. In contrast, suprathreshold or threshold-level stimulation, which induces action potentials, has been found to regulate heart-brain interaction [15, 25]. Our results extend these findings by showing that the intensity parameter is critically affected by the stimulation site, revealing a more complex interaction. Notably, at certain stimulation sites (e.g., F3 lateral, posterior, and medial), HBC increases significantly at approximately suprathreshold or threshold-level stimulation, whereas others (e.g., F3) respond at lower intensities. This finding challenges the conventional assumption that one global stimulation threshold, i.e., the motor threshold, suffices to individualize stimulation across the cortex. Instead, this divergence in effective stimulation intensities across targets suggests that the functional threshold for DLPFC rTMS differs from the standard MT, underscoring the need for HBC-based markers to guide individualized dosing [39].

For most targets, the repeatability of HBC remained low across three sessions, even at sites with significant intensity effects (F3, F3 medial and posterior). Only F3 anterior and F3 lateral at higher intensities showed consistent repeatability (Fig. 3a). Interestingly, sessions 2 and 3 were more consistent than comparisons involving session 1 (Fig. 3b). This may reflect the novelty effect of the first session, which reduces in subsequent sessions. Similar observations emerge in studies reporting an initial heart rate deceleration that disappears at follow-up after 30 days of rTMS [44]. These findings highlight the importance of considering session-specific factors and adaptation effects in heart rate dynamics.

Beyond cortical stimulation, TMS can activate sub-scalp nerves and muscles, potentially causing site-specific pain. Since pain is known to affect heart rate, we monitored and quantified pain and side effects to distinguish TMS-related from those related to pain and related side effects. Although both factors influenced HBC, we identified a TMS-induced change in HBC that persisted beyond those confounding effects. Mild pain is a common TMS side effect arising from directly activating nerves and muscles beneath the scalp. This nociceptive stimulation triggers the autonomic nervous system and alters heart rate [45]. Additional side effects, such as facial twitches [46], sensory alterations [47], and fatigue [48] can affect both the sympathetic and parasympathetic pathways, influencing HBC. Emotional responses like fear can heighten sympathetic activity and elevate heart rate [49], yet paradoxically induce transient bradycardia via vagal activation [50]. These bidirectional effects underscore the intricate relationship between autonomic regulation and TMS-induced side effects, likely driving the non-linear impact of TMS on HBC.

Across F3-surrounding targets, F3 lateral produced the largest and most reproducible increase in HBC with a clear intensity dependence. Crucially, this pattern persisted after modeling side effects. In the single-side-effect GAM (Table S5), F3 lateral showed no significant interactions with muscle contraction, scalp pain, or headache, while retaining a strong target main effect and a significant non-linear %MT×target interaction. These findings indicate that the robust HBC modulation at F3 lateral is unlikely to arise from non-specific somatosensory confounds and instead reflects genuine neurophysiological engagement. These statistical observations are consistent with a connectivity-based mechanism. Resting-state fMRI work has shown that left DLPFC subregions that are more negatively correlated with the sgACC tend to yield superior antidepressant outcomes [51]. Direct coordinate-wise comparisons further indicate that clinically superior DLPFC targets are slightly more anterior and lateral and exhibit stronger sgACC anticorrelation than their less effective counterparts with the 5-cm rule method versus the non-responder site (approximately at MNI coordinate –46, +23, +49 vs. –41, +17, +55) [52]. In this context, F3 lateral approximates an antero-lateral DLPFC sector that is more strongly embedded in an sgACC-anticorrelated network than the canonical EEG F3 position [13], providing a network-level rationale for its side-effect-resistant, intensity-dependent HBC effects.

Conceptually, HBC might offer a physiological read-out of frontal–vagal network engagement. The observation that prefrontal TMS systematically modulates heart-linked dynamics supports the broader view that brain stimulation can probe, and potentially normalize, dysregulated brain–autonomic interactions relevant to affect regulation and arousal control. This positions HBC as a candidate target-engagement biomarker to complement symptom measures in personalization frameworks.

Several limitations of our study warrant careful consideration. First, our experiments were conducted in healthy participants and evaluated only acute physiological endpoints; thus, generalizability to clinical populations and relevance to long-term symptom change remain uncertain. Second, we did not acquire participant-specific resting-state fMRI, precluding subject-level estimates of DLPFC–sgACC connectivity. Third, intensity selection was based on rMT, which does not equalize the prefrontal cortical E-field, leaving potential within-target, between-target and between-participant variability [53]; E-field-informed dosing could mitigate this and improve reliability. Finally, although the apparent advantage of the F3 lateral target persisted after statistically controlling for common side effects, residual somatosensory or state-dependent confounds cannot be fully excluded.

Future work should integrate subject-specific connectivity with detailed electric-field modeling to test two hypotheses: 1) the F3 lateral site preferentially engages the sgACC-anticorrelated DLPFC subnetwork; and 2) such engagement mediates both physiological effects (e.g., HBC modulation) and clinical improvement. Clinical evaluation should then proceed to randomized trials comparing HBC-guided targeting with connectivity–guided and standard Beam-F3/5-cm approaches, with pre-specified mediation analyses. Standardization of acquisition pipelines and E-field reporting will be important for reproducibility and eventual clinical translation.

In summary, this study substantially adds to previous discoveries by revealing that the NCG-TMS 2.0 protocol can robustly influence cardiac activity [21]. Advancing beyond prior research [38], we uncover a non-linear modulation of HBC dependent on stimulation intensity and specific targets within the left DLPFC. Notably, at the lateral F3 site, intensity-dependent stimulation reliably enhances HBC compared to sham. By evaluating these effects across multiple sessions within the same participants, we demonstrated high variability in HBC changes for most stimulation targets, but also the robustness of our findings for specific sites, particularly anterior and lateral F3. Furthermore, we explore the directionality of TMS-induced modulation of HBC, showing a consistent decrease in heart rate only at the F3 lateral target with suprathreshold stimulation. Future investigations should distinguish TMS-induced electric fields from neural circuits, particularly sgACC–DLPFC connectivity, to clarify their autonomic impact. These insights could deepen our understanding of the heart-brain axis and inform targeted therapeutic strategies.

## Data availability

The full dataset is made available at https://osf.io/8qpv3/. All computational steps and related code are available at https://gitlab.gwdg.de/tms-localization/papers/tms-hbc.

## CRediT authorship contribution statement

**Zi-jian Feng:** Data curation, Formal analysis, Investigation, Methodology, Project administration, Validation, Writing – original draft, Writing – review & editing. **Sandra Martin:** Formal analysis, Investigation, Methodology, Validation, Visualization, Writing – original draft, Writing – review & editing. **Ole Numssen:** Conceptualization, Investigation, Methodology, Project administration, Validation, Writing – original draft, Writing – review & editing. **Konstantin Weise:** Conceptualization, Investigation, Project administration, Writing – review & editing. **Ying Jing:** Investigation, Project administration. **Gerasimos Gerardos:** Data curation, Investigation. **Carla Martin:** Data curation, Investigation. **Gesa Hartwigsen:** Conceptualization, Funding acquisition, Investigation, Project administration, Resources, Supervision, Writing – review & editing. **Thomas Knösche:** Conceptualization, Funding acquisition, Investigation, Project administration, Resources, Supervision, Writing – review & editing.

## Acknowledgments

We would like to express our sincere gratitude to Ariana Novak, Benjamin Kalloch, Jeremy Ranke, and Nico Leng for their invaluable contributions to the data collection process. We are also deeply thankful to Philipp Kuhnke for his insightful guidance and advice during the data analysis phase. Their efforts and expertise were integral to the success of this study. This work was supported by Lise Meitner Excellence funding from the Max Planck Society, the European Research Council (ERC-2021-COG 101043747), and the German Research Foundation (HA 6314/3-1, HA 6314/4-2, HA 6314/9-1). Zijian Feng was supported by the Deqing Hospital of Hangzhou Normal University and Medical and Health Research Project of Zhejiang Province (2023XY042). Ying Jing was supported by the China Scholarship Council. Ole Numssen and Konstantin Weise were supported by the Federal Ministry of Education Germany (Bundesministerium für Bildung und Forschung, BMBF, Grant no. 01GQ2201 to Thomas Knösche).

## References

1. Lefaucheur JP, Aleman A, Baeken C, Benninger DH, Brunelin J, Di Lazzaro V et al. Evidence-based guidelines on the therapeutic use of repetitive transcranial magnetic stimulation (rTMS): An update (2014-2018). Clin Neurophysiol. 2020; 131: 474–528. 10.1016/j.clinph.2019.11.002.

2. Rossini PM, Burke D, Chen R, Cohen LG, Daskalakis Z, Di Iorio R et al. Non-invasive electrical and magnetic stimulation of the brain, spinal cord, roots and peripheral nerves: Basic principles and procedures for routine clinical and research application. An updated report from an I.F.C.N. Committee. Clin Neurophysiol. 2015; 126: 1071–1107. 10.1016/j.clinph.2015.02.001.

3. Numssen O, van der Burght CL, Hartwigsen G. Revisiting the focality of non-invasive brain stimulation - Implications for studies of human cognition. Neurosci Biobehav Rev. 2023; 149: 105154. 10.1016/j.neubiorev.2023.105154.

4. Martin S, Frieling R, Saur D, Hartwigsen G. TMS over the pre-SMA enhances semantic cognition via remote network effects on task-based activity and connectivity. Brain Stimul. 2023; 16: 1346–1357. 10.1016/j.brs.2023.09.009.

5. Feng ZJ, Deng XP, Zhao N, Jin J, Yue J, Hu YS et al. Resting-State fMRI Functional Connectivity Strength Predicts Local Activity Change in the Dorsal Cingulate Cortex: A Multi-Target Focused rTMS Study. Cereb Cortex. 2022; 32: 2773–2784. 10.1093/cercor/bhab380.

6. Spellman T, Liston C. Toward Circuit Mechanisms of Pathophysiology in Depression. Am J Psychiatry. 2020; 177: 381–390. 10.1176/appi.ajp.2020.20030280.

7. Berger S, Kliem A, Yeragani V, Bar KJ. Cardio-respiratory coupling in untreated patients with major depression. J Affect Disord. 2012; 139: 166–171. 10.1016/j.jad.2012.01.035.

8. Koenig J, Kemp AH, Beauchaine TP, Thayer JF, Kaess M. Depression and resting state heart rate variability in children and adolescents - A systematic review and meta-analysis. Clin Psychol Rev. 2016; 46: 136–150. 10.1016/j.cpr.2016.04.013.

9. Schmausser M, Raab M, Laborde S. The dynamic role of the left dlPFC in neurovisceral integration: Differential effects of theta burst stimulation on vagally mediated heart rate variability and cognitive-affective processing. Psychophysiology. 2024; 61: e14606. 10.1111/psyp.14606.

10. Thayer JF, Lane RD. Claude Bernard and the heart-brain connection: further elaboration of a model of neurovisceral integration. Neurosci Biobehav Rev. 2009; 33: 81–88. 10.1016/j.neubiorev.2008.08.004.

11. Thayer JF, Ahs F, Fredrikson M, Sollers JJ, 3rd, Wager TD. A meta-analysis of heart rate variability and neuroimaging studies: implications for heart rate variability as a marker of stress and health. Neurosci Biobehav Rev. 2012; 36: 747–756. 10.1016/j.neubiorev.2011.11.009.

12. Iseger TA, van Bueren NER, Kenemans JL, Gevirtz R, Arns M. A frontal-vagal network theory for Major Depressive Disorder: Implications for optimizing neuromodulation techniques. Brain Stimul. 2020; 13: 1–9. 10.1016/j.brs.2019.10.006.

13. Fox MD, Buckner RL, White MP, Greicius MD, Pascual-Leone A. Efficacy of transcranial magnetic stimulation targets for depression is related to intrinsic functional connectivity with the subgenual cingulate. Biol Psychiatry. 2012; 72: 595–603. 10.1016/j.biopsych.2012.04.028.

14. Cole EJ, Phillips AL, Bentzley BS, Stimpson KH, Nejad R, Barmak F et al. Stanford Neuromodulation Therapy (SNT): A Double-Blind Randomized Controlled Trial. Am J Psychiatry. 2022; 179: 132–141. 10.1176/appi.ajp.2021.20101429.

15. Iseger TA, Padberg F, Kenemans JL, Gevirtz R, Arns M. Neuro-Cardiac-Guided TMS (NCG-TMS): Probing DLPFC-sgACC-vagus nerve connectivity using heart rate - First results. Brain Stimul. 2017; 10: 1006–1008. 10.1016/j.brs.2017.05.002.

16. Jiao Y, Cheng C, Jia M, Chu Z, Song X, Zhang M et al. Neuro-cardiac-guided transcranial magnetic stimulation: Unveiling the modulatory effects of low-frequency and high-frequency stimulation on heart rate. Psychophysiology. 2024; 61: e14631. 10.1111/psyp.14631.

17. Cheng C, Jiao Y, Jia M, Jin J, Peng X, Zhang M et al. Neuro-cardiac coupling predicts heart rate regulation effects of neuro-cardiac-guided transcranial magnetic stimulation (TMS). Brain Stimul. 2024; 17: 1134–1136. 10.1016/j.brs.2024.09.007.

18. Kaur M, Michael JA, Hoy KE, Fitzgibbon BM, Ross MS, Iseger TA et al. Investigating high- and low-frequency neuro-cardiac-guided TMS for probing the frontal vagal pathway. Brain Stimul. 2020; 13: 931–938. 10.1016/j.brs.2020.03.002.

19. Zwienenberg L, Iseger TA, Dijkstra E, Rouwhorst R, van Dijk H, Sack AT et al. Neuro-cardiac guided rTMS as a stratifying method between the ‘5cm’ and ‘BeamF3’ stimulation clusters. Brain Stimul. 2021; 14: 1070–1072. 10.1016/j.brs.2021.07.005.

20. Thut G, Veniero D, Romei V, Miniussi C, Schyns P, Gross J. Rhythmic TMS causes local entrainment of natural oscillatory signatures. Curr Biol. 2011; 21: 1176–1185. 10.1016/j.cub.2011.05.049.

21. Dijkstra E, van Dijk H, Vila-Rodriguez F, Zwienenberg L, Rouwhorst R, Coetzee JP et al. Transcranial Magnetic Stimulation-Induced Heart-Brain Coupling: Implications for Site Selection and Frontal Thresholding-Preliminary Findings. Biol Psychiatry Glob Open Sci. 2023; 3: 939–947. 10.1016/j.bpsgos.2023.01.003.

22. Rouwhorst R, van Oostrom I, Dijkstra E, Zwienenberg L, van Dijk H, Arns M. Vasovagal syncope as a specific side effect of DLPFC-rTMS: A frontal-vagal dose-finding study. Brain Stimul. 2022; 15: 1233–1235. 10.1016/j.brs.2022.08.015.

23. Heck K, Ferguson J, Sege C, Huffman S, Brown T, George MS. The clinical efficacy of heart rate as a biomarker for TMS. Transcranial Magnetic Stimulation. 2025; 4. 10.1016/j.transm.2025.100180.

24. Alario AA, Pace BD, Niciu MJ, Trapp NT. Transcranial magnetic stimulation induces heart rate decelerations independent of treatment outcome. Brain Stimul. 2023; 16: 1044–1046. 10.1016/j.brs.2023.06.005.

25. Iseger TA, Padberg F, Kenemans JL, van Dijk H, Arns M. Neuro-Cardiac-Guided TMS (NCG TMS): A replication and extension study. Biol Psychol. 2021; 162: 108097. 10.1016/j.biopsycho.2021.108097.

26. Iseger TA, Arns M, Downar J, Blumberger DM, Daskalakis ZJ, Vila-Rodriguez F. Cardiovascular differences between sham and active iTBS related to treatment response in MDD. Brain Stimul. 2020; 13: 167–174. 10.1016/j.brs.2019.09.016.

27. Rossi S, Antal A, Bestmann S, Bikson M, Brewer C, Brockmoller J et al. Safety and recommendations for TMS use in healthy subjects and patient populations, with updates on training, ethical and regulatory issues: Expert Guidelines. Clin Neurophysiol. 2021; 132: 269–306. 10.1016/j.clinph.2020.10.003.

28. Mir-Moghtadaei A, Caballero R, Fried P, Fox MD, Lee K, Giacobbe P et al. Concordance Between BeamF3 and MRI-neuronavigated Target Sites for Repetitive Transcranial Magnetic Stimulation of the Left Dorsolateral Prefrontal Cortex. Brain Stimul. 2015; 8: 965–973. 10.1016/j.brs.2015.05.008.

29. Pascual-Leone A, Rubio B, Pallardo F, Catala MD. Rapid-rate transcranial magnetic stimulation of left dorsolateral prefrontal cortex in drug-resistant depression. Lancet. 1996; 348: 233–237. 10.1016/s0140-6736(96)01219-6.

30. Polomano RC, Galloway KT, Kent ML, Brandon-Edwards H, Kwon KN, Morales C et al. Psychometric Testing of the Defense and Veterans Pain Rating Scale (DVPRS): A New Pain Scale for Military Population. Pain Med. 2016; 17: 1505–1519. 10.1093/pm/pnw105.

31. Giustiniani A, Vallesi A, Oliveri M, Tarantino V, Ambrosini E, Bortoletto M et al. A questionnaire to collect unintended effects of transcranial magnetic stimulation: A consensus based approach. Clin Neurophysiol. 2022; 141: 101–108. 10.1016/j.clinph.2022.06.008.

32. sjPlot: Data Visualization for Statistics in Social Science. R package version 2.8.17. https://CRAN.R-project.org/package=sjPlot, 2024, Accessed Date Accessed 2024 Accessed.

33. Lüdecke D, Ben-Shachar M, Patil I, Waggoner P, Makowski D. performance: An R Package for Assessment, Comparison and Testing of Statistical Models. Journal of Open Source Software. 2021; 6. 10.21105/joss.03139.

34. Makowski D, Ben-Shachar M, Patil I, Lüdecke D. Methods and Algorithms for Correlation Analysis in R. Journal of Open Source Software. 2020; 5: 2306. 10.21105/joss.02306.

35. Lüdecke D. ggeffects: Tidy Data Frames of Marginal Effects from Regression Models. Journal of Open Source Software. 2018; 3: 772. 10.21105/joss.00772.

36. Wood SN. Generalized additive models: an introduction with R. (chapman and hall/CRC, 2017).

37. Bates D, Mächler M, Bolker B, Walker S. Fitting Linear Mixed-Effects Models Usinglme4. Journal of Statistical Software. 2015; 67: 1–48. 10.18637/jss.v067.i01.

38. Dijkstra ESA, Frandsen SB, van Dijk H, Duecker F, Taylor JJ, Sack AT et al. Probing prefrontal-sgACC connectivity using TMS-induced heart–brain coupling. Nature Mental Health. 2024; 2: 809–817. 10.1038/s44220-024-00248-8.

39. Numssen O, Kuhnke P, Weise K, Hartwigsen G. Electric-field-based dosing for TMS. Imaging Neurosci (Camb). 2024; 2: 1–12. 10.1162/imag_a_00106.

40. Hartwigsen G, Silvanto J. Noninvasive Brain Stimulation: Multiple Effects on Cognition. Neuroscientist. 2023; 29: 639–653. 10.1177/10738584221113806.

41. Tang AD, Bennett W, Bindoff AD, Bolland S, Collins J, Langley RC et al. Subthreshold repetitive transcranial magnetic stimulation drives structural synaptic plasticity in the young and aged motor cortex. Brain Stimul. 2021; 14: 1498–1507. 10.1016/j.brs.2021.10.001.

42. Kuhnke P, Meyer L, Friederici AD, Hartwigsen G. Left posterior inferior frontal gyrus is causally involved in reordering during sentence processing. Neuroimage. 2017; 148: 254–263. 10.1016/j.neuroimage.2017.01.013.

43. Crewther BT, Kasprzycka W, Cook CJ, Rola R. Impact of one HF-rTMS session over the DLPFC and motor cortex on acute hormone dynamics and emotional state in healthy adults: a sham-controlled pilot study. Neurol Sci. 2022; 43: 651–659. 10.1007/s10072-021-05335-7.

44. Alario AA, Pace BD, Niciu MJ, Trapp NT. Transcranial magnetic stimulation-associated heart rate decelerations attenuate after a TMS treatment course for depression. Brain Stimul. 2024; 17: 1155–1156. 10.1016/j.brs.2024.09.011.

45. Porges SW. Cardiac vagal tone: a physiological index of stress. Neurosci Biobehav Rev. 1995; 19: 225–233. 10.1016/0149-7634(94)00066-a.

46. Proske U, Gandevia SC. The proprioceptive senses: their roles in signaling body shape, body position and movement, and muscle force. Physiol Rev. 2012; 92: 1651–1697. 10.1152/physrev.00048.2011.

47. Yang W, Chen T, He R, Goossens R, Huysmans T. Autonomic responses to pressure sensitivity of head, face and neck: Heart rate and skin conductance. Appl Ergon. 2024; 114: 104126. 10.1016/j.apergo.2023.104126.

48. Matuz A, van der Linden D, Kisander Z, Hernadi I, Kazmer K, Csatho A. Enhanced cardiac vagal tone in mental fatigue: Analysis of heart rate variability in Time-on-Task, recovery, and reactivity. PLoS One. 2021; 16: e0238670. 10.1371/journal.pone.0238670.

49. Messina G, Monda A, Messina A, Di Maio G, Monda V, Limone P et al. Relationship between Non-Invasive Brain Stimulation and Autonomic Nervous System. Biomedicines. 2024; 12. 10.3390/biomedicines12050972.

50. Forte G, Troisi G, Pazzaglia M, Pascalis V, Casagrande M. Heart Rate Variability and Pain: A Systematic Review. Brain Sci. 2022; 12. 10.3390/brainsci12020153.

51. Cash RFH, Zalesky A. Personalized and Circuit-Based Transcranial Magnetic Stimulation: Evidence, Controversies, and Opportunities. Biol Psychiatry. 2024; 95: 510–522. 10.1016/j.biopsych.2023.11.013.

52. Herbsman T, Avery D, Ramsey D, Holtzheimer P, Wadjik C, Hardaway F et al. More lateral and anterior prefrontal coil location is associated with better repetitive transcranial magnetic stimulation antidepressant response. Biol Psychiatry. 2009; 66: 509–515. 10.1016/j.biopsych.2009.04.034.

53. Numssen O, Martin S, Williams K, Knosche TR, Hartwigsen G. Quantification of subject motion during TMS via pulsewise coil displacement. Brain Stimul. 2024; 17: 1045–1047. 10.1016/j.brs.2024.08.009.

