## Supplementary Methods for "Target-Specificity and Repeatability in Neuro-Cardiac-Guided TMS for Heart-Brain Coupling"

### Supplementary Materials

#### Supplementary Methods

##### Power Analysis

Our within-subject factorial design includes three factors: Target, Intensity, and Session. This resulted in 7 (Target) * 16 (Intensity) * 3 (Session) = 336 measurements per participant. Based on an anticipated small effect size of 0.1, with alpha set to 0.05 and statistical power to 0.8, the minimum required sample size was calculated to be 16 participants (https://aaroncaldwell.us/SuperpowerBook/). To account for an estimated 20% dropout rate due to technical difficulties or participant discomfort (e.g., pain), we invited 20 participants to complete all three sessions.

##### MRI Acquisition

For each participant, MRI scans were acquired using a 3T MRI scanner (Siemens Verio or Skyra fit) equipped with a 32-channel head coil to obtain high-resolution T1-weighted images. The scanning parameters were as follows: T1: MPRAGE ADNI with 176 sagittal slices; repetition time (TR): 2.3 s; echo time (TE): 2.98 ms; field of view (FoV): 256 mm; voxel size: 1 x 1 x 1 mm; no slice gap; flip angle: 9°. If high-quality scans (less than two years old) were available in the in-house database, they were used instead of new acquisitions.

##### TMS Setup

For TMS, we used a MagPro X100 stimulator (MagVenture, Farum, Denmark) with a passively cooled MCF-B65 figure-of-eight coil. Sham stimulation was applied using the corresponding sham coil (MCF-P-B65), which generates the same acoustic noise as the active coil but reduces the magnetic field strength by approximately 80%. Participants were blinded to the type of stimulation (active vs. sham). Coil positioning was guided by a neuro-navigation system (TMS Navigator, Localite, Sankt Augustin, Germany) with a Polaris Spectra camera (NDI, Waterloo, Canada), and coil placements were recorded for each TMS pulse to ensure correct placement throughout the experiment.

At the beginning of the first session, the resting motor threshold (rMT) for the first dorsal interosseous (FDI) muscle was manually determined as the minimum stimulator intensity required to elicit motor evoked potentials with an amplitude of at least 50 μV in 5 out of 10 consecutive TMS pulses, following the relative frequency method [1]. Electromyography (EMG) signals from the right FDI muscle were used to detect motor evoked potentials during the rMT determination procedure.

#### ECG recording

Electrocardiogram (ECG) data were recorded during stimulation using the REFA8 68-channel amplifier system (TMSi, Oldenzaal, The Netherlands) with Brain Vision Recorder software (Brain Vision, MedCaT B.V.). A visual inspection of the ECG trace was conducted to ensure signal quality before the experiment began. Prior to testing, participants were seated and instructed to remain relaxed, minimize movement, and refrain from speaking. ECG electrodes were placed as follows: the ground electrode above the left breast, the reference electrode above the right breast, and the active electrode below the left breast. ECG signals were continuously recorded during the stimulation protocol.

#### Side effects measurement

To assess participants' tolerance, a single 10 Hz TMS train at 120% rMT, the maximum stimulation intensity of the protocol, was applied to each target before rTMS administration. Unpleasantness levels were assessed using the Discrete Visual Pain Rating Scale (DVPRS) [2]. Pain scores for each target are shown in Figure S1. If pain scores exceeded 6 (out of 10) points (indicating a level above 'moderate'), the coil angle was adjusted from 45° to 90° to reduce discomfort. This adjustment was necessary for six participants. Despite these modifications, five targets from three participants were ultimately excluded due to persistent pain scores exceeding 6 points. Sham stimulation was performed at one of the five targets, F3 or one of the four surrounding targets, randomly assigned for each participant. Sessions 2 and 3 consisted of repetitions of both active and sham NCG-TMS 2.0 protocols conducted during Session 1. The target order was randomized across participants, and the same order was maintained across all sessions.

After each session, participants completed a stimulation-related side-effects checklist (Table S1) assessing 20 symptoms: scalp pain, toothache, scalp tingling, peripheral-nerve tingling, itching, burning/heat, headache, tinnitus/noise, skin sensations, unintended muscle contraction, fatigue, sleepiness, hearing changes, mood changes (depression; euphoria), nausea, neck stiffness/pain, coil pressure, anxiety/nervousness, difficulty concentrating, and “other (specify).” For each symptom they reported: (i) severity (0–4: 0 = none, 1 = mild, 2 = moderate, 3 = considerable, 4 = strong); (ii) onset (beginning / middle / toward the end / after the end of stimulation); (iii) duration (stopped quickly; stopped during the train; stopped at the end; or continued after the end with duration in minutes); and (iv) location (e.g., head/arm/finger; diffuse vs. localized near the stimulation site) [3]. For the statistical modeling, we calculated a sum score of reported side effects per session and target. This was entered into statistical models as a predictor next to the pain rating from the DVPRS scale. Supplementary tables and figures display the frequency of pain ratings and side effects: Figure S1 (DVPRS), Figure S2 and S3 (side effects), and Table S2 (frequency of side effects).

#### Individual Heart-Brain Coupling

The ECG data analysis in this study was adapted from the original analyses [4] (https://github.com/brainclinics/HBC). Initially, ECG signals were processed using a bidirectional fourth-order Butterworth bandpass filter with a frequency range of 0.5 to 20 Hz, effectively removing low-frequency noise and high-frequency artifacts. The R peak, a prominent upward deflection in the QRS complex of the ECG signal that represents ventricular depolarization, was identified using the 'signal.findpeaks' function from SciPy [5] along with the ‘ecg_peaks’ function from NeuroKit2 toolbox [6]. Manual correction was subsequently applied to improve detection accuracy. Signals requiring manual corrections for more than 10% of the total R peaks for any target were excluded from the analysis, and one target from a participant was excluded due to insufficient usable signal.

Following R peak detection, the time differences between two consecutive R-waves in the ECG (RR intervals) were corrected for ectopic beats, and the HR was computed for each RR interval. To smooth the HR data, a moving average of five consecutive HR values was applied, followed by interpolation across all ECG time points. A 1.5-second Hann window was applied to reduce abrupt fluctuations and enhance overall data consistency.

Time-frequency analysis was conducted using the 'tfr_array_morlet' function from MNE-Python [7]. This generated a time-frequency representation of the ECG signal across a frequency range of 0.02 to 0.18 Hz, with increments of 0.0025 Hz. High temporal resolution (3 cycles) and high frequency resolution (10 cycles) allowed for a detailed examination of low-frequency oscillations. Data padding was applied at both the start and end of the sequence to facilitate the analysis of these low-frequency components.

HBC was quantified by the power at 0.0625 Hz corresponding to one TMS cycle comprising 5 seconds of stimulation and 11 seconds of rest. The analysis was applied to all 26 blocks of ECG data. However, only the 16 blocks corresponding to the NCG-TMS 2.0 protocol were retained, while the initial 8 resting blocks and the last 2 resting blocks were excluded to mitigate edge effects. All analyses were conducted using the raw values of HBC power. The high frequency resolution HBC value for each block was then calculated and used for subsequent analyses. Plots for each target across all three sessions for each participant are provided in figure S4.

**Mixed-Effects Regression Analysis**

To analyze the relationship between HBC power of each block and our experimental factors (stimulation intensity and target location) as well as nuisance variables (pain and other side effects), we used mixed-effects regression. Plotting the raw data per target and session suggested a non-linear relationship between stimulation intensity and HBC power (Fig. S5). To account for this nonlinearity, we set up a generalized additive model (GAM). GAMs determine the optimal shape of the curve fit (i.e., the degree of *smoothness*) for non-linear responses via penalized regression. This data-driven adjustment became necessary as the observed non-linearity violated the assumptions of the pre-registered ANOVA approach, making GAMs a more appropriate choice for modeling these effects. The GAM was fitted with a Gamma distribution and a log link function to account for the non-normal distribution of HBC power. This transformation was determined as appropriate based on distribution diagnostics using the package fitdistrplus [8] to ensure residual normality and homoscedasticity. We included both fixed and random effects in the GAM to appropriately model the data structure and dependencies. The model contained main effects for *session*, *target*, and percentage intensity of MT (*%MT*) as well as the interaction of *target* and *%MT*. We further included two covariates: (i) per participant and target assessed *pain scores* to account for a potential interdependence of HBC with higher pain sensations, and (ii) a sum score of the per participant, session, and target measured *side effects*, which summarize sensations such as headache, nausea, and skin sensation. Moreover, we included the interaction of *%MT* and *pain*, aiming to identify a potential modulation of HBC through intensity-dependent pain values.

The splines of the continuous predictors *%MT*, *pain*, and *side effects* were fitted with thin-plate regression splines. The interaction of *%MT* and *target* was fitted with by-target individual splines, resulting in individual curves for each level of *target*. For the interaction of *%MT* and *pain* a tensor product interaction term with a cubic regression spline was used to accommodate the differing scales of both variables. The number of basis dimensions for each smoothing spline was checked via model diagnostics available in *mgcv* after the first model set-up and appropriately updated to reach $p > 0.05$ to avoid overfitting. Random effects included random intercepts for *participant* and *block*, and by-participant random slopes for *session* (EQ 1).

$$
\begin{aligned}
\textit{HBC Power} \sim \quad & \textit{Session} + \\
& \textit{Stimulation Target} + \\
& \textit{\%MT} + \\
& \textit{Pain Score} + \\
& \textit{Side Effects} + \\
& \textit{Stimulation Target : \%MT} \\
& \textit{Pain : \%MT} + \\
& \textit{(1 + Session | Participant)} + \\
& \textit{(1 | Block)}
\end{aligned}
\qquad \text{(EQ 1)}
$$

We also fitted a standard linear regression model to confirm our results without relaxing the linearity assumption. To this end, a generalized linear mixed-effects model (GLMM) with a Gamma distribution and a log link function was set up with the same fixed and random effects as the GAM (see EQ 1).

In all regression analyses, we employed simple contrast coding for categorical factors and mean-centering for continuous predictors to ensure interpretability and comparability of the results. Additionally, we applied a false discovery rate (FDR) correction to the p-values obtained from the model summaries to control for multiple comparisons. Analyses were carried out in R version 4.4.0 (R Core Team, 2023) using the packages lme4, mgcv, modelbased, ggeffects, performance, and sjPlot [9-14].

**Repeatability Analysis**

To assess the repeatability of block-level measurements across multiple sessions for each target, we calculated the Intraclass Correlation Coefficient (ICC) using a two-way random-effects model. This model assumes that both targets and sessions were selected at random, which allows for generalization beyond the specific samples studied. Specifically, we assessed the consistency of HBC power values across three sessions for each block within a target. The ICC formula used is as follows:

$$ICC = \frac{MSb - MSw}{MSb + (k-1)MSw} \qquad (EQ\ 2)$$

MSb (Mean Square Between Subjects) quantifies the variability in measurements between participants, averaged across all sessions, MSw (Mean Square Within Subjects) reflects the variability within individual participants across sessions, and $k$ represents the number of sessions (here: $k = 3$).

The ICC was computed using the Pingouin Python package [15], which employs an ANOVA-based approach to estimate the variance components. This analysis was performed to determine the repeatability of block-level measurements across repeated sessions, thereby providing insight into the stability and consistency of TMS-induced HBC responses.

High ICC values indicate that most of the observed variability is driven by between-participant differences, suggesting good repeatability. Conversely, lower ICC values imply substantial variability within participants across sessions, potentially undermining the repeatability of these measurements. In addition, we calculated pairwise ICC values between each session to further explore the consistency between individual sessions.

**Analyses of Heart Rate Modulation Direction**

Previous studies have demonstrated that vagus nerve stimulation elicits a rapid autonomic response within the same cardiac cycle, with effects typically lasting only 1–2 heartbeats [16]. Similarly, pupil size, a well-established indicator of autonomic nervous system activity, has been shown to respond to TMS within a 2-second window [17]. Drawing on these findings, the HR analysis was conducted using a 2-second post-stimulation window to capture potential rapid autonomic responses. For pre-train HR, two reference intervals were employed: 1) Baseline, which excluded data from the first two blocks of each session to ensure HR stability; 2) Pre-train, defined as the 2-second window immediately preceding stimulation. If two consecutive R peaks could not be identified within the defined 2-second window for HR calculation, HR was instead determined using the first two consecutive R peaks identified either before or after the stimulation period.

### Supplementary Figures

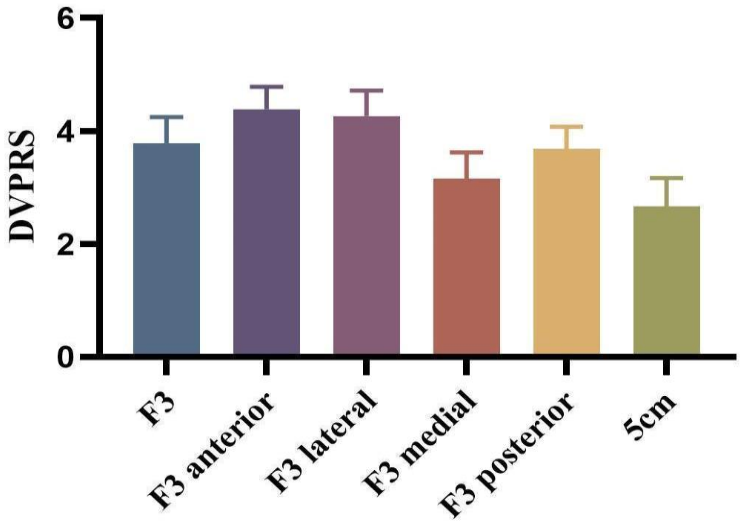

**Figure S1. Defense and Veterans Pain Rating Scale (DVPRS) Scores for Different Stimulation Targets.** Error bars represent the standard error of the mean (SEM)

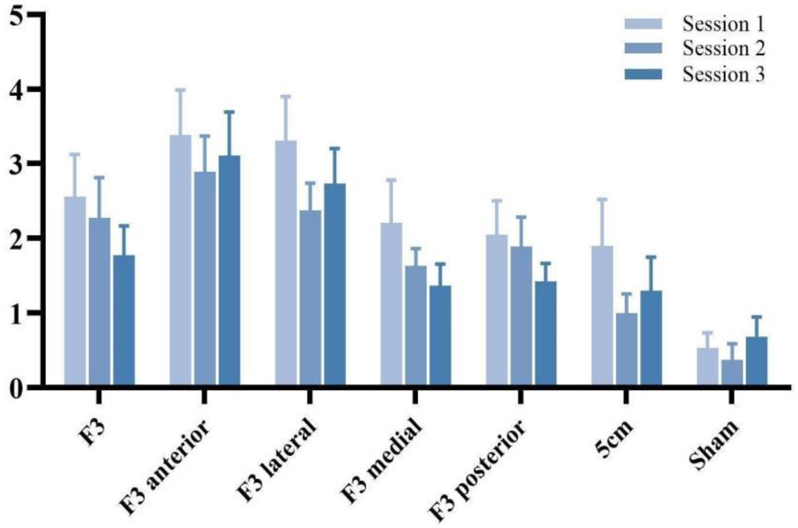

**Figure S2. Side Effect Scores across Different Stimulation Targets.** Total side effect scores across three sessions for different stimulation targets, as measured by a questionnaire assessing potential side effects (Giustiniani et al., 2022). The y-axis represents the total side effect score, while the x-axis shows various stimulation targets. Error bars indicate the standard error of the mean (SEM).

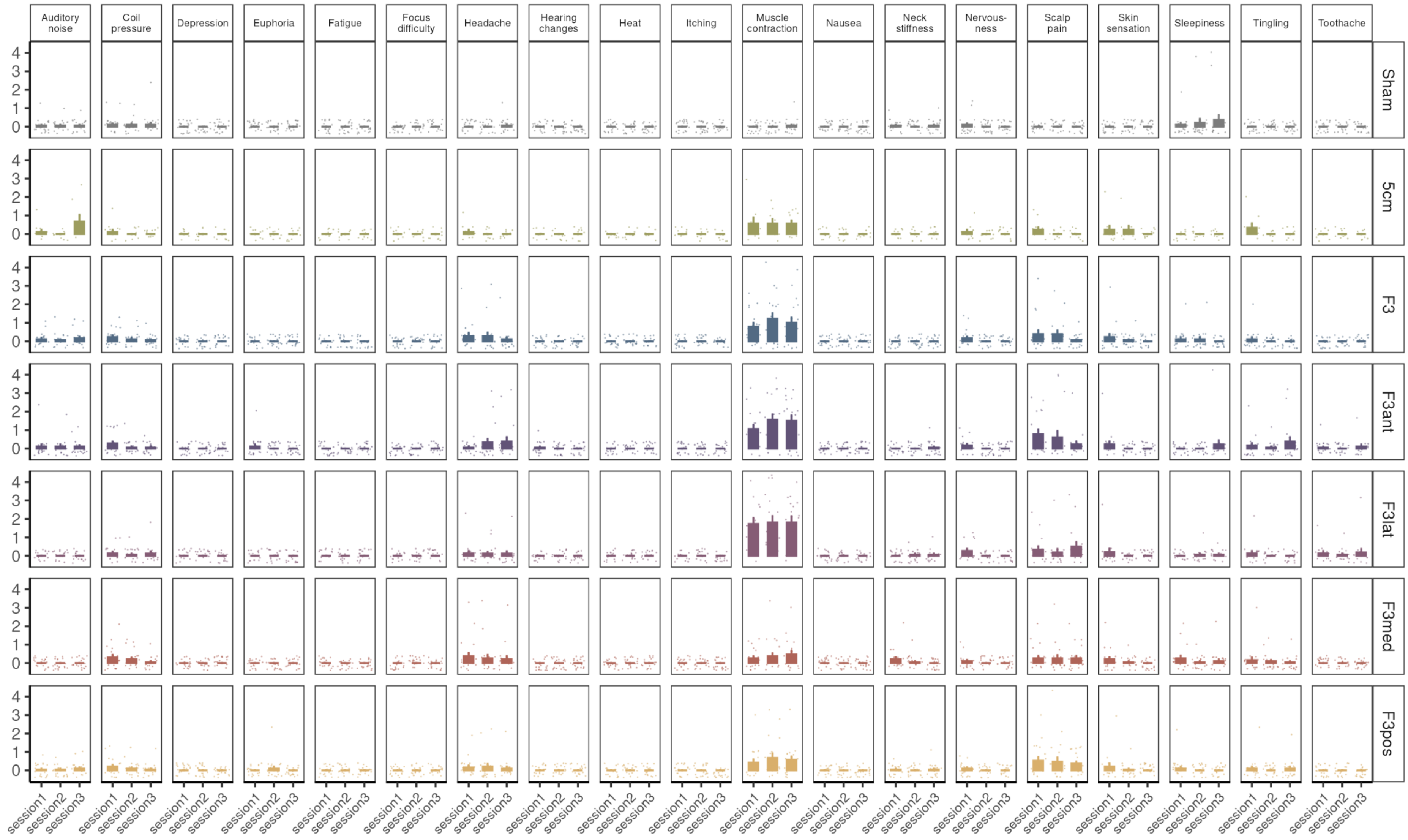

**Figure S3. Individual side effects across targets and sessions.** Each side effect was rated on a scale from 0 to 4 after each target stimulation in each session. Bar plots display mean scores with standard error and individual data points plotted on top.

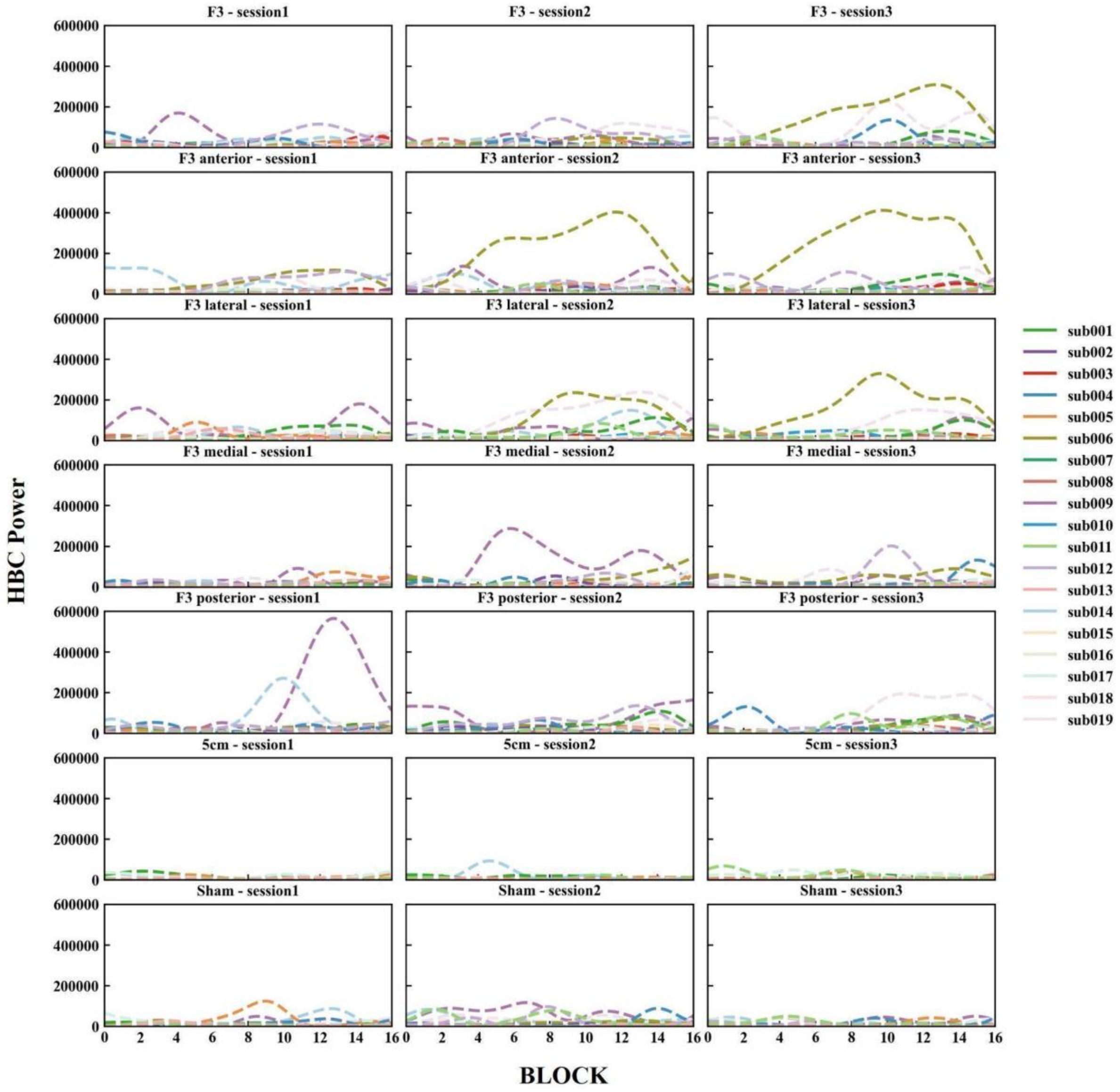

**Figure S4. Individual Raw HBC Power across Stimulation Targets and Sessions.** Raw heart-brain coupling (HBC) power values for individual subjects across different stimulation targets and sessions. Each panel represents a specific stimulation target across three sessions. The y-axis shows HBC power, while the x-axis represents block numbers, corresponding to increasing TMS stimulation intensities. Different colored dashed lines indicate individual subject data.

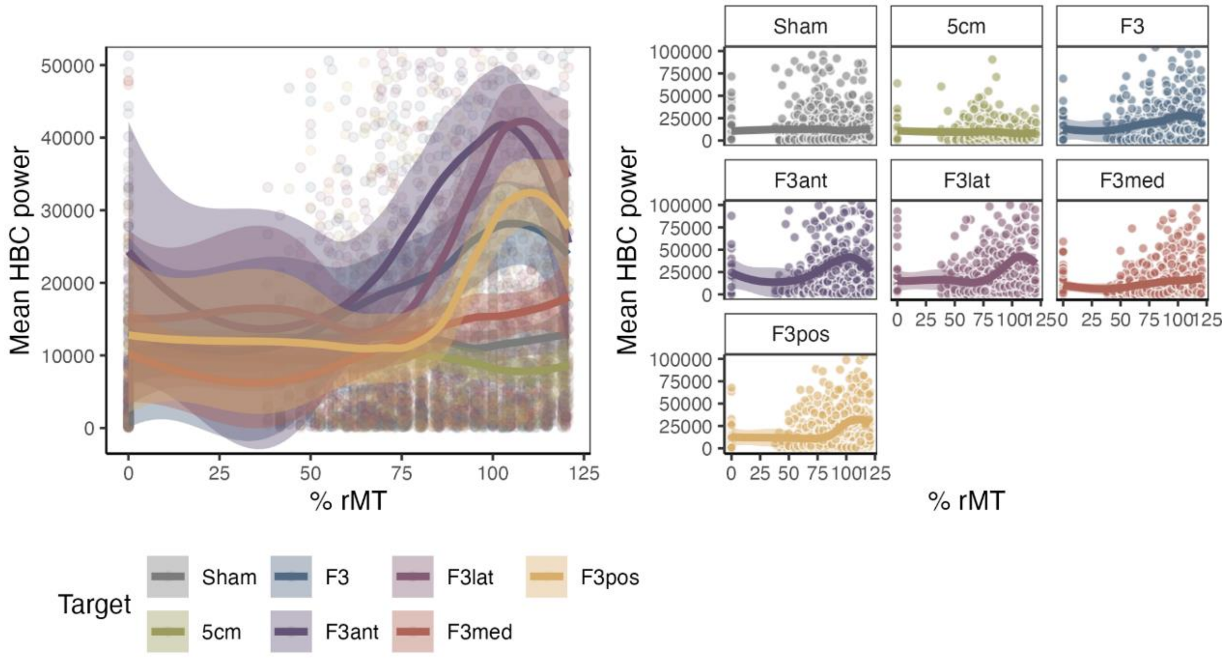

**Figure S5. Raw HBC data.** The left panel shows raw data with smoothed lines (method = Loess) per target on top. The right panel shows raw data with smooth lines with facets per target.

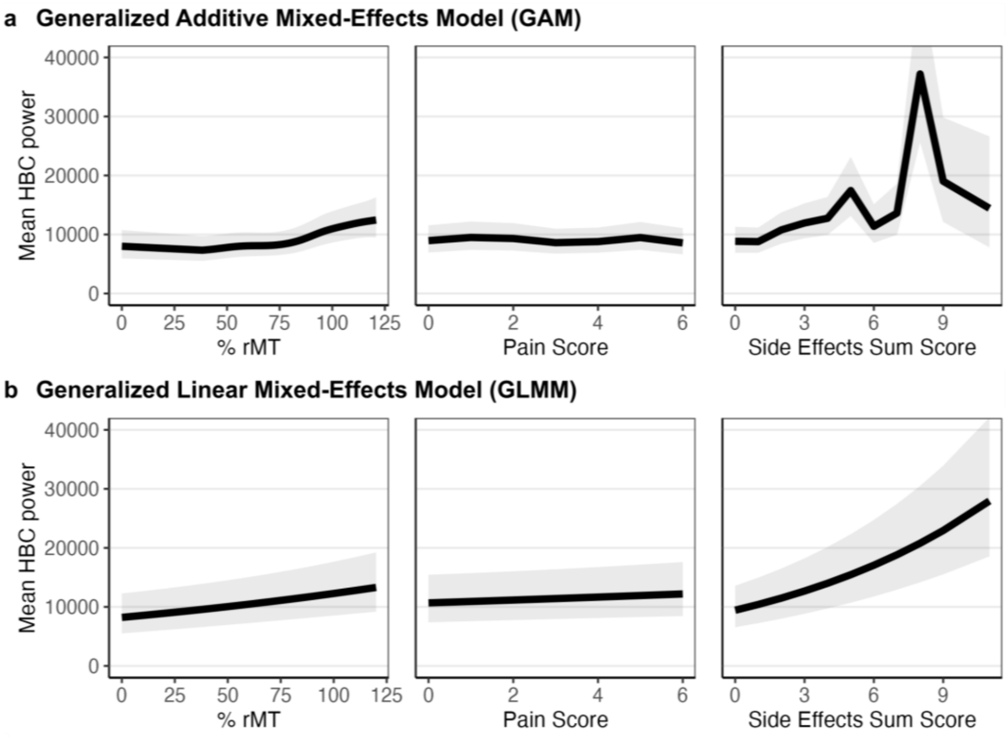

**Figure S6. Estimated Marginal Predictions for %MT, Pain, and Side Effects from the Generalized Additive Regression Model.** Panel a shows smoothing splines with confidence intervals from the GAM for the main effects of %MT, pain, and side effects. Only side effects had a significant effect on HBC (see Table 1). Panel b shows estimated regression lines with confidence intervals from the GLMM. In this model, both %rMT and side effects were significant predictors (see Table 1).

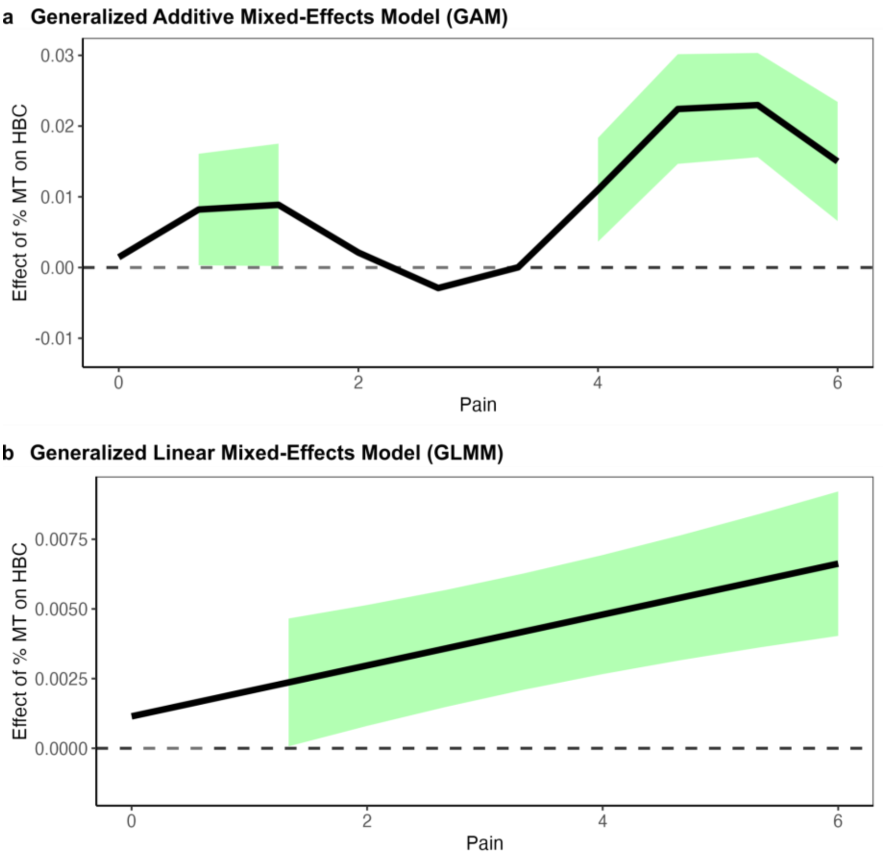

**Figure S7. Interaction of % rMT and Pain Scores.** The figure shows the results of the simple slopes analysis for the interaction between %MT and Pain in the (a) generalized additive model and (b) generalized linear regression model.

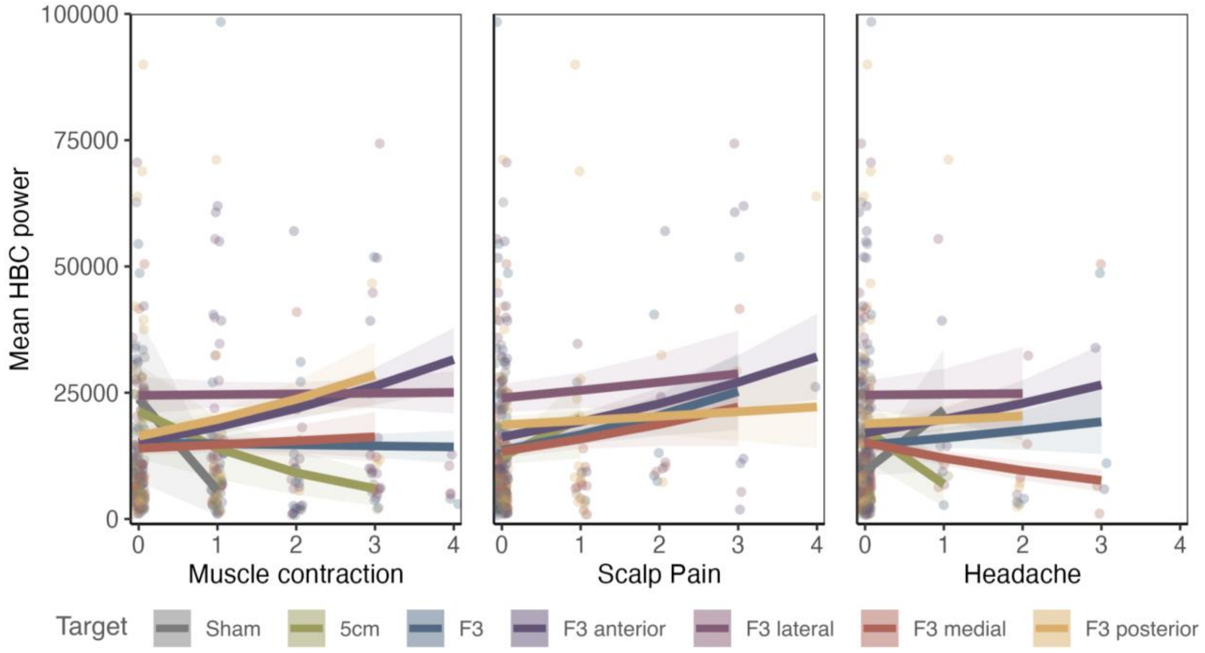

**Fig. S8. Interaction of three most common side effects (muscle contraction, scalp pain, and headache) and Stimulation Targets.** The figure shows the average marginal mean for each target at observed values of each side effect in the data. Subject-, session, and target-wise raw data are plotted below.

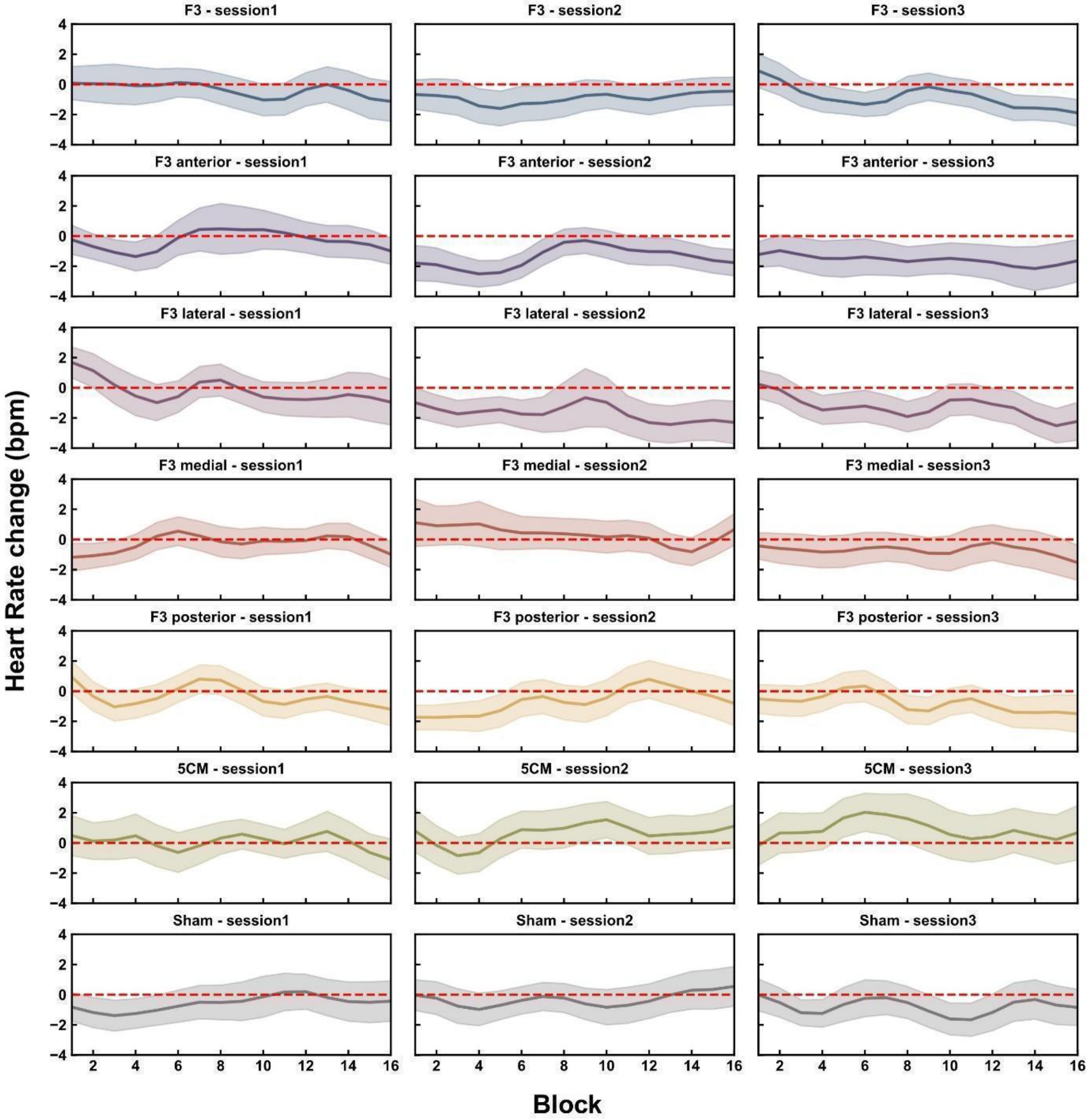

**Figure S9. Heart Rate (HR) Changes (in beats per minute, bpm) Post-train Compared to Baseline across Stimulation Targets for each Session.** Shaded areas indicate the standard error of the mean (SEM). The red dashed line marks zero change, serving as a reference for HR variation.

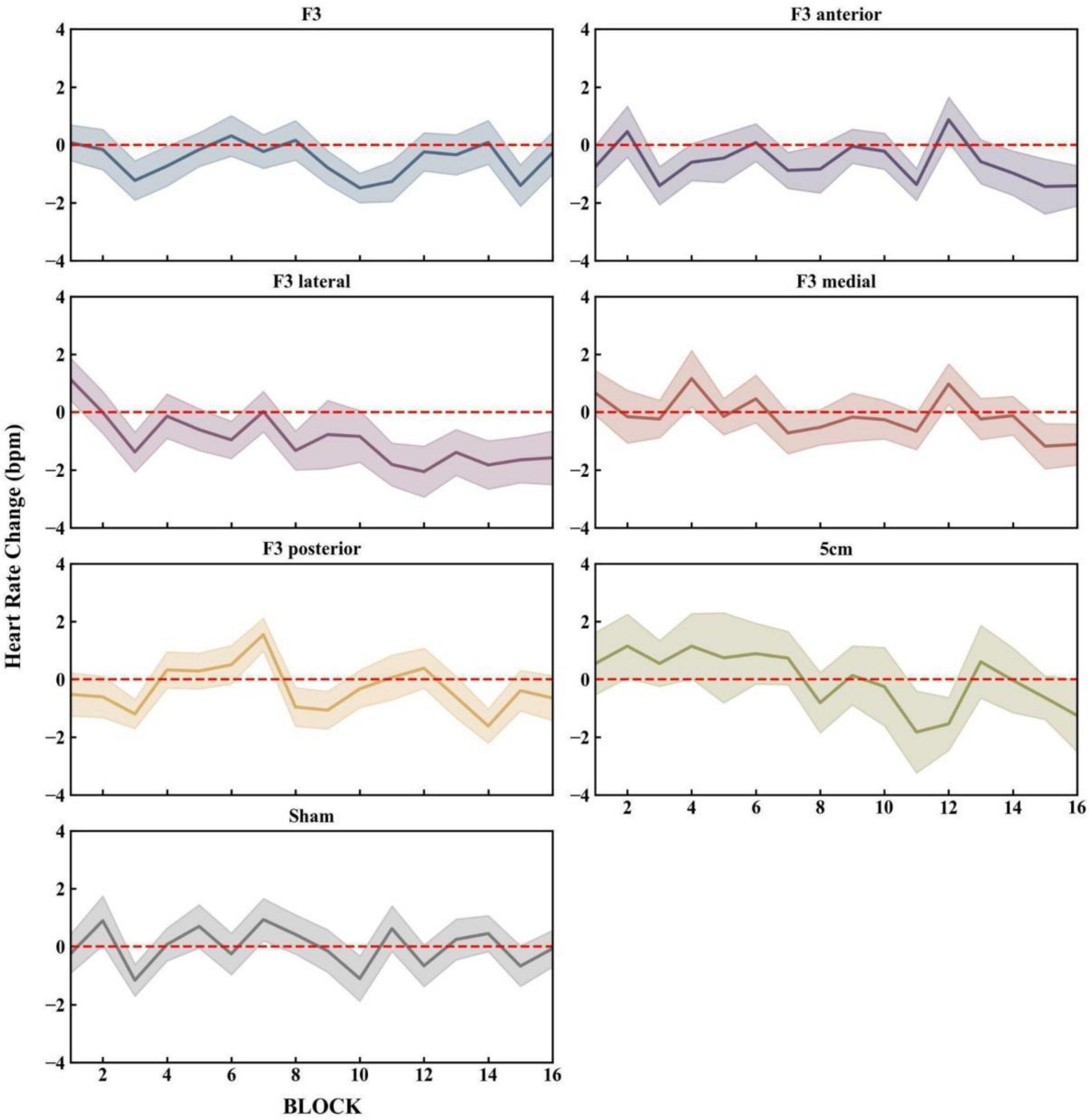

**Figure S10. Heart Rate (HR) Changes (in beats per minute, bpm) Post-train Compared to Pre-train across Stimulation Targets.** Shaded areas indicate the standard error of the mean (SEM). The red dashed line marks zero change, serving as a reference for HR variation.

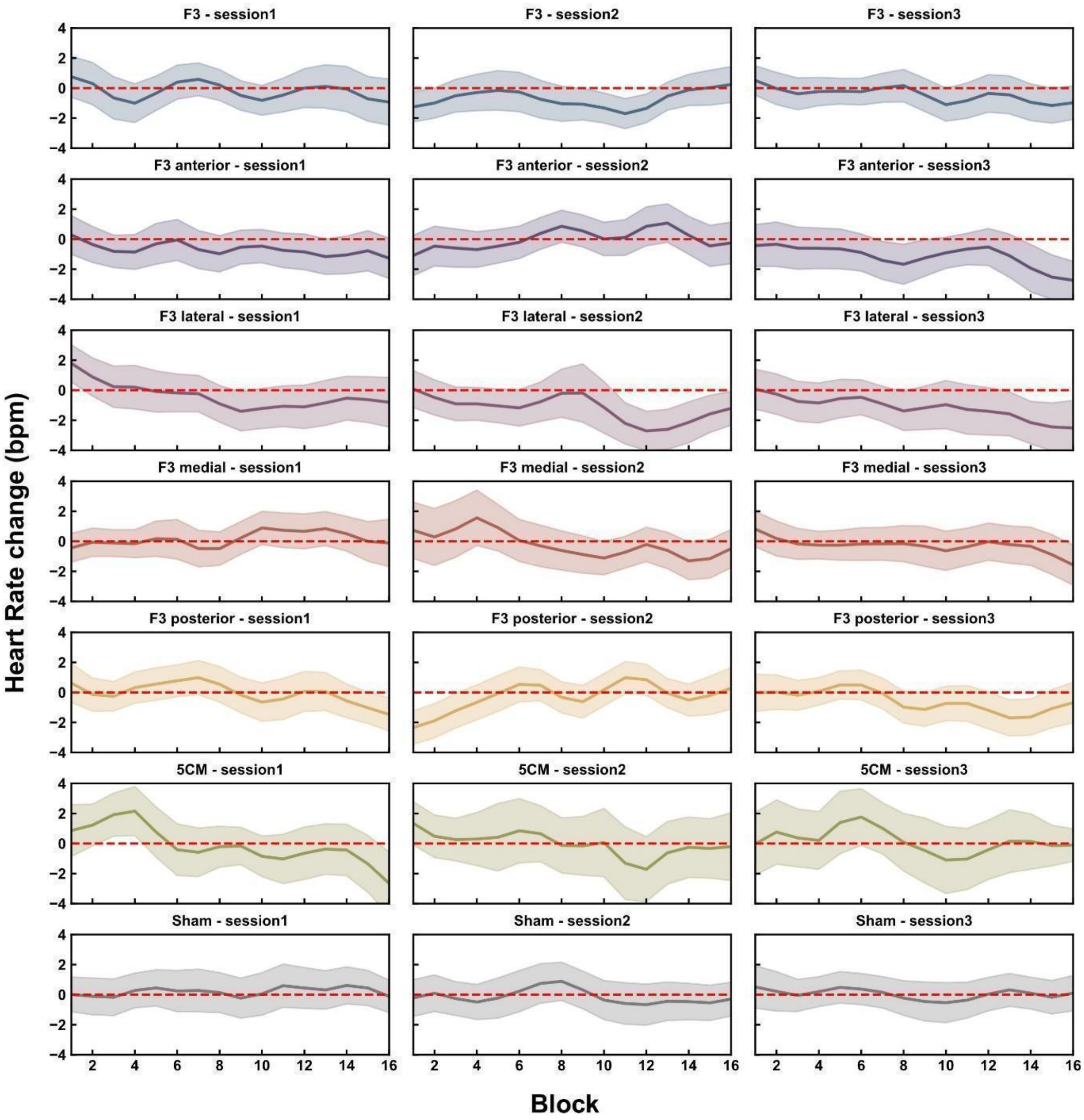

**Figure S11. Heart Rate (HR) Changes (in beats per minute, bpm) Post-train Compared to Pre-train across Stimulation Targets across Sessions.** Shaded areas indicate the standard error of the mean (SEM). The red dashed line marks zero change, serving as a reference for HR variation.

### Supplementary Tables

**Table S1. Stimulation related sensations**

| | When did the sensations begin? | | | | | How long did it last? | | | | Location |
|---|---|---|---|---|---|---|---|---|---|---|
| | Degree (from 0 to 4) | At the beginning of the stimulation | In the middle of the stimulation | Towards the end of the stimulation | After the end of the stimulation | It stopped quickly | It stopped in the middle of the stimulatio n | It stopped at the end of the stimulatio n | it stopped after the end of the stimulatio n (duration in min) | (e.g., head, arm, finger) Diffuse/ localized/ close to stimulation |
| **1. Scalp pain** | | | | | | | | | | |
| **2. Toothache** | | | | | | | | | | |
| **3. Tingling at scalp** | | | | | | | | | | |
| **4. Tingling (peripheral nerves)** | | | | | | | | | | |
| **5. Itching** | | | | | | | | | | |
| **6. Burning or heat** | | | | | | | | | | |
| **7. Headache** | | | | | | | | | | |
| **8. Noise (e.g., tinnitus)** | | | | | | | | | | |
| **9. Skin sensation** | | | | | | | | | | |
| **10. Muscle contraction (excluding “targeted” MEPs)** | | | | | | | | | | |
| **11. Fatigue** | | | | | | | | | | |
| **12. Sleepiness** | | | | | | | | | | |
| **13. Hearing changes** | | | | | | | | | | |
| **14. Mood changes (depression)** | | | | | | | | | | |
| **15. Mood changes (euphoria)** | | | | | | | | | | |
| **16. Nausea** | | | | | | | | | | |
| **17. Neck stiffness/pain** | | | | | | | | | | |
| **18. Coil pressure** | | | | | | | | | | |

| |
|---|
| **19. Anxiety/ Nervousness** |
| **20. Difficulty in concentrating** |
| **21. Other (specify)** |

*Note. To be filled after each session. Did you experience any of the following sensations? Please answer by inserting the number that corresponds to the degree of the experienced discomfort, with 0 (None), 1 (Mild), 2 (Moderate), 3 (Considerable), 4 (Strong). Please, specify when the sensation started and how long it lasted.*

**Table S2. Sum of individual side effects across sessions.**

| Side effect | Sum across participants and sessions |
|---|---|
| Muscle contraction | 281 |
| Scalp pain | 103 |
| Headache | 59 |
| Coil pressure | 44 |
| Sleepiness | 33 |
| Tingling | 28 |
| Noise | 26 |
| Skin sensation | 26 |
| Nervousness | 17 |
| Neck stiffness | 12 |
| Toothache | 9 |
| Mood changes euphoria | 4 |
| Hearing changes | 1 |
| Burning/heat | 0 |
| Difficulty concentrating | 0 |
| Fatigue | 0 |
| Itching | 0 |
| Mood changes depression | 0 |
| Nausea | 0 |

**Table S3. Results from the Generalized Linear Mixed-effects Model (GLMM).**

| | HBC Power | | | |
|---|---|---|---|---|
| ***Predictors*** | ***Estimate*** | ***std. Error*** | ***Statistic*** | ***p*** |
| Session [Session 2] | 1.15 | 0.14 | 1.16 | 3.36e-01 |
| Session [Session 3] | 1.03 | 0.18 | 0.18 | 9.06e-01 |
| Target [5cm] | 0.90 | 0.06 | -1.61 | 1.72e-01 |
| **Target [F3]** | 0.86 | 0.06 | -2.31 | **3.95e-02** |
| Target [F3 anterior] | 1.07 | 0.08 | 0.94 | 4.26e-01 |
| **Target [F3 lateral]** | 1.33 | 0.10 | 3.93 | **5.55e-04** |
| **Target [F3 medial]** | 0.84 | 0.05 | -2.87 | **1.11e-02** |
| Target [F3 posterior] | 1.09 | 0.07 | 1.33 | 2.70e-01 |
| %MT | 1.00 | 0.00 | 3.77 | 9.06e-01 |
| Pain | 1.02 | 0.01 | 1.89 | 1.03e-01 |
| **Side Effects** | 1.10 | 0.01 | 9.07 | **1.48e-18** |
| %MT x Target [5cm] | 1.00 | 0.00 | 0.03 | 9.75e-01 |
| %MT x Target [F3] | 1.00 | 0.00 | 0.92 | 4.26e-01 |
| **%MT x Target [F3 ant]** | 1.01 | 0.00 | 2.60 | **2.24e-02** |
| **%MT x Target [F3 lat]** | 1.01 | 0.00 | 3.82 | **6.33e-04** |
| **%MT x Target [F3 med]** | 1.00 | 0.00 | 2.48 | **2.81e-02** |
| **%MT x Target [F3 pos]** | 1.01 | 0.00 | 3.20 | **4.41e-03** |
| **%MT x Pain** | 1.00 | 0.00 | 3.35 | **3.10e-03** |
| Marginal /Conditional $R^2$ | 0.069 / 0.560 | | | |

*Note.* Significant effects are marked in bold. Note that no separate term is reported for the interaction of %MT and the sham target, since sham was modeled as baseline in the simple contrast coding scheme of the factor target. Parameter estimates are exponentiated coefficients and represent ratios on the response scale. Reported p-values are FDR-corrected.

**Table S4. Output of Model Comparisons.**

| Model Comparisons | | | | | | |
|---|---|---|---|---|---|---|
| ***Name*** | ***AIC*** | ***AICc*** | ***BIC*** | ***R2*** | ***RMSE*** | ***Sigma*** |
| gamm_1 | 116683.05 | 116687.33 | 117402.83 | 0.49 | 26072.01 | 1.03 |
| gamm_MT | 116729.61 | 116733.26 | 117393.85 | 0.48 | 26518.37 | 1.04 |
| gamm_Pain | 116767.88 | 116771.98 | 117472.36 | 0.47 | 26803.66 | 1.05 |

*Note*. Models were compared using the function compare_performance() from the performance package (Lüdecke et al., 2021). Gamm_1 is the reported model including both Pain and %MT as predictors (see equation 1). We compared this model to models without each predictor and found that including both predictors significantly improved the model.

**Table S5. Results from the Generalized Additive Model (GAM) with three strongest side effects.**

| | Mean HBC Power | | | |
|---|---|---|---|---|
| ***Predictors*** | ***Estimate / edf*** | ***std. Error*** | ***Statistic*** | ***p*** |
| Session [Session 2] | 1.09 | 0.31 | 0.32 | 8.08e-01 |
| Session [Session 3] | 1.01 | 0.29 | 0.04 | 9.72e-01 |
| Target [5cm] | 2.27 | 0.55 | 3.38 | **1.42e-03** |
| Target [F3] | 2.72 | 0.63 | 4.32 | **3.83e-05** |
| Target [F3 ant] | 3.16 | 0.74 | 4.93 | **2.72e-06** |
| Target [F3 lat] | 4.47 | 01.05 | 6.38 | **9.72e-10** |
| Target [F3 med] | 2.68 | 0.62 | 4.23 | **5.48e-05** |
| Target [F3 pos] | 3.49 | 0.81 | 5.41 | **2.76e-07** |
| Muscle contr. x Target [Sham] | 0.24 | 0.07 | -5.06 | **1.49e-06** |
| Muscle contr. x Target [5cm] | 0.66 | 0.05 | -5.70 | **5.86e-08** |
| Muscle contr. x Target [F3] | 0.99 | 0.03 | -0.40 | 7.63e-01 |
| Muscle contr. x Target [F3 ant] | 1.20 | 0.04 | 5.34 | **3.59e-07** |
| Muscle contr. x Target [F3 lat] | 1.01 | 0.03 | 0.18 | 9.03e-01 |
| Muscle contr. x Target [F3 med] | 1.05 | 0.06 | 0.89 | 4.76e-01 |
| Muscle contr. x Target [F3 pos] | 1.20 | 0.05 | 4.10 | **8.93e-05** |
| Scalp pain x Target [Sham] | 1.00 | 0.00 | NaN | NaN |
| Scalp pain x Target [5cm] | 1.61 | 0.34 | 2.28 | **3.57e-02** |
| Scalp pain x Target [F3] | 1.23 | 0.07 | 3.78 | **3.31e-04** |
| Scalp pain x Target [F3 ant] | 1.19 | 0.04 | 4.66 | **9.59e-06** |
| Scalp pain x Target [F3 lat] | 1.06 | 0.05 | 1.20 | 3.16e-01 |
| Scalp pain x Target [F3 med] | 1.18 | 0.08 | 2.63 | **1.39e-02** |
| Scalp pain x Target [F3 pos] | 1.04 | 0.05 | 0.86 | 4.87e-01 |
| Headache x Target [Sham] | 2.28 | 0.64 | 2.93 | **6.03e-03** |
| Headache x Target [5cm] | 0.39 | 0.11 | -3.28 | **1.93e-03** |
| Headache x Target [F3] | 1.10 | 0.07 | 1.54 | 1.75e-01 |
| Headache x Target [F3 ant] | 1.16 | 0.06 | 2.74 | **1.06e-02** |
| Headache x Target [F3 lat] | 1.01 | 0.10 | 0.06 | 9.72e-01 |
| Headache x Target [F3 med] | 0.80 | 0.04 | -4.46 | **2.31e-05** |
| Headache x Target [F3 pos] | 1.04 | 0.08 | 0.53 | 7.02e-01 |
| s(%MT) | 0.0009 | | 0.00 | 3.68e-01 |
| s(Pain) | 0.79 | | 6.54 | 5.77e-02 |
| s(%MT x Target [Sham]) | 0.0004 | | 0.00 | 7.63e-01 |
| s(%MT x Target [5cm]) | 1.38 | | 0.27 | **4.05e-02** |
| s(%MT x Target [F3]) | 2.51 | | 1.23 | **3.03e-05** |
| s(%MT x Target [F3 ant]) | 2.96 | | 2.20 | **0.00e+00** |
| s(%MT x Target [F3 lat]) | 3.53 | | 3.90 | **0.00e+00** |
| s(%MT x Target [F3 med]) | 5.49 | | 2.26 | **0.00e+00** |
| s(%MT x Target [F3 pos]) | 2.87 | | 3.55 | **0.00e+00** |
| ti(%MT x Pain) | 1.28 | | 3.11 | **0.00e+00** |
| Marginal / Conditional $R^2$ | 0.453 | | | |

*Note.* Significant effects are marked in bold. Parameter estimates are exponentiated coefficients and represent ratios on the response scale. For smooths and smooth-by-factor interactions, the effective degrees of freedom (edf) are shown instead of parameter estimates and standard errors. Reported p-values are FDR-corrected.

### References

1. Rossini PM, Burke D, Chen R, Cohen LG, Daskalakis Z, Di Iorio R *et al.* Non-invasive electrical and magnetic stimulation of the brain, spinal cord, roots and peripheral nerves: Basic principles and procedures for routine clinical and research application. An updated report from an I.F.C.N. Committee. Clin Neurophysiol. 2015; 126: 1071-1107. https://doi.org/10.1016/j.clinph.2015.02.001.

2. Polomano RC, Galloway KT, Kent ML, Brandon-Edwards H, Kwon KN, Morales C *et al.* Psychometric Testing of the Defense and Veterans Pain Rating Scale (DVPRS): A New Pain Scale for Military Population. Pain Med. 2016; 17: 1505-1519. https://doi.org/10.1093/pm/pnw105.

3. Giustiniani A, Vallesi A, Oliveri M, Tarantino V, Ambrosini E, Bortoletto M *et al.* A questionnaire to collect unintended effects of transcranial magnetic stimulation: A consensus based approach. Clin Neurophysiol. 2022; 141: 101-108. https://doi.org/10.1016/j.clinph.2022.06.008.

4. Dijkstra E, van Dijk H, Vila-Rodriguez F, Zwienenberg L, Rouwhorst R, Coetzee JP *et al.* Transcranial Magnetic Stimulation-Induced Heart-Brain Coupling: Implications for Site Selection and Frontal Thresholding-Preliminary Findings. Biol Psychiatry Glob Open Sci. 2023; 3: 939-947. https://doi.org/10.1016/j.bpsgos.2023.01.003.

5. Virtanen P, Gommers R, Oliphant TE, Haberland M, Reddy T, Cournapeau D *et al.* SciPy 1.0: fundamental algorithms for scientific computing in Python. Nat Methods. 2020; 17: 261-272. https://doi.org/10.1038/s41592-019-0686-2.

6. Makowski D, Pham T, Lau ZJ, Brammer JC, Lespinasse F, Pham H *et al.* NeuroKit2: A Python toolbox for neurophysiological signal processing. Behav Res Methods. 2021; 53: 1689-1696. https://doi.org/10.3758/s13428-020-01516-y.

7. Gramfort A, Luessi M, Larson E, Engemann DA, Strohmeier D, Brodbeck C *et al.* MEG and EEG data analysis with MNE-Python. Front Neurosci. 2013; 7: 267. https://doi.org/10.3389/fnins.2013.00267.

8. Delignette-Muller ML, Dutang C. fitdistrplus: An R Package for Fitting Distributions. Journal of Statistical Software. 2015; 64: 1 - 34. https://doi.org/10.18637/jss.v064.i04.

9. sjPlot: Data Visualization for Statistics in Social Science. R package version 2.8.17. https://CRAN.R-project.org/package=sjPlot, 2024, Accessed Date Accessed 2024 Accessed.

10. Lüdecke D, Ben-Shachar MS, Patil I, Waggoner P, Makowski D. performance: An R package for assessment, comparison and testing of statistical models. Journal of Open Source Software. 2021; 6. https://doi.org/10.21105/joss.03139.

11. Makowski D, Ben-Shachar MS, Patil I, Lüdecke D. Methods and algorithms for correlation analysis in R. Journal of Open Source Software. 2020; 5: 2306. https://doi.org/10.21105/joss.02306.

12. Lüdecke D. ggeffects: Tidy data frames of marginal effects from regression models. Journal of Open Source Software. 2018; 3: 772. https://doi.org/10.21105/joss.00772.

13. Wood SN. Generalized additive models: an introduction with R. (chapman and hall/CRC, 2017).

14. Bates D, Mächler M, Bolker B, Walker S. Fitting Linear Mixed-Effects Models Using lme4. Journal of Statistical Software. 2015; 67: 1 - 48. https://doi.org/10.18637/jss.v067.i01.

15. Vallat R. Pingouin: statistics in Python. Journal of Open Source Software. 2018; 3: 1026. https://doi.org/10.21105/joss.01026.

16. Iseger TA, van Bueren NER, Kenemans JL, Gevirtz R, Arns M. A frontal-vagal network theory for Major Depressive Disorder: Implications for optimizing neuromodulation techniques. Brain Stimul. 2020; 13: 1-9. https://doi.org/10.1016/j.brs.2019.10.006.

17. Niehaus L, Guldin B, Meyer B. Influence of transcranial magnetic stimulation on pupil size. J Neurol Sci. 2001; 182: 123-128. https://doi.org/10.1016/s0022-510x(00)00462-7.
